# Microbes on the Mind: Multi-modal Neuroimaging Reveals Gut-Brain Axes in Infants

**DOI:** 10.64898/2026.09.23.753825

**Authors:** Tengfei Li, Yue Yang, Seo-Ho Cho, Brittany R. Howell, Zhengwang Wu, Weiyan Yin, Jed T. Elison, Khoi Minh Huynh, Sahar Ahmad, Pew-Thian Yap, Gang Li, Li Wang, Hongtu Zhu, Weili Lin, UNC/UMN Baby Connectome Project Consortium

## Abstract

The gut microbiome undergoes rapid maturation after birth and has been associated with cognition and mental health, yet its relation to early brain development remains poorly defined. We analyzed 223 typically developing children aged 0–3 years with multimodal MRI and paired fecal shotgun metagenomics. Microbiome-wide association analyses identified associations with cortical morphology, white-matter microstructure, and functional connectivity. Ridge-regularized canonical correlation analysis identified two principal covariance patterns: a structural–metabolic mode and a functional–taxonomic mode. Sparse canonical partial least squares recovered the principal CCA score and phenotype patterns within the same sample. Longitudinal canonical-score slopes were not significantly correlated; in autoregressive cross-lag models, however, neuroimaging CCA1 scores predicted microbiome CCA1 scores 1–6 months later. These results indicate cross-domain covariance and temporal ordering within the measured windows but do not establish causality.

## Introduction

The microbiome–gut–brain axis encompasses reciprocal signaling between the gastrointestinal tract and central nervous system through immune, neuroendocrine, and vagal pathways [1–3]. Gut microbes and enterochromaffin cells can influence neurotransmitter availability, including serotonin, and modulate innate and adaptive immunity [4, 5]. Conversely, central autonomic and hormonal pathways can alter gastrointestinal physiology and microbial composition [6]. Perturbations of this system have been associated with irritable-bowel syndrome, obesity, and several neuropsychiatric disorders [6].

Early life is a period of rapid microbial colonization and brain maturation, but evidence that these processes covary remains limited. Infant studies using 16S rRNA sequencing or shotgun metagenomics have reported associations of microbial diversity with regional brain volume, resting-state connectivity, and functional near-infrared signals [7–10]. Interpretation has been limited by small samples, low taxonomic resolution, restricted metabolic characterization, and incomplete imaging coverage.

To examine these relationships, we analyzed an accelerated longitudinal cohort of typically developing children from birth to three years. Fecal samples underwent shotgun metagenomics, and structural, diffusion, and resting-state functional MRI characterized cortical morphology, white-matter microstructure, and functional connectivity. We used regularized canonical correlation analysis to estimate multivariate microbiome–brain covariance and sparse canonical partial least squares to determine whether the principal CCA score, feature, and phenotype patterns were recoverable under a different multivariate objective. Participant-level resampling quantified held-out stability.

## Results

### Study cohort

The BCP-Enriched sample comprised a subset of the UNC/UMN Baby Connectome Project [11]. We collected 955 fecal samples from 305 children aged 0–3 years. Matching stool and MRI observations within ±30 days yielded 489 paired sessions from 223 children. After imaging quality control, the analytic datasets included 424 structural MRI sessions from 200 children, 176 diffusion MRI sessions from 120 children, and 400 resting-state fMRI sessions from 196 children; 107 children contributed all three modalities. Table 1 summarizes participant characteristics. Figure 1A–D summarizes longitudinal sampling and modality-specific availability, and Figure 1E outlines the microbiome and neuroimaging preprocessing, phenotype extraction, developmental, and cross-domain analytic workflow.

**Figure 1.**
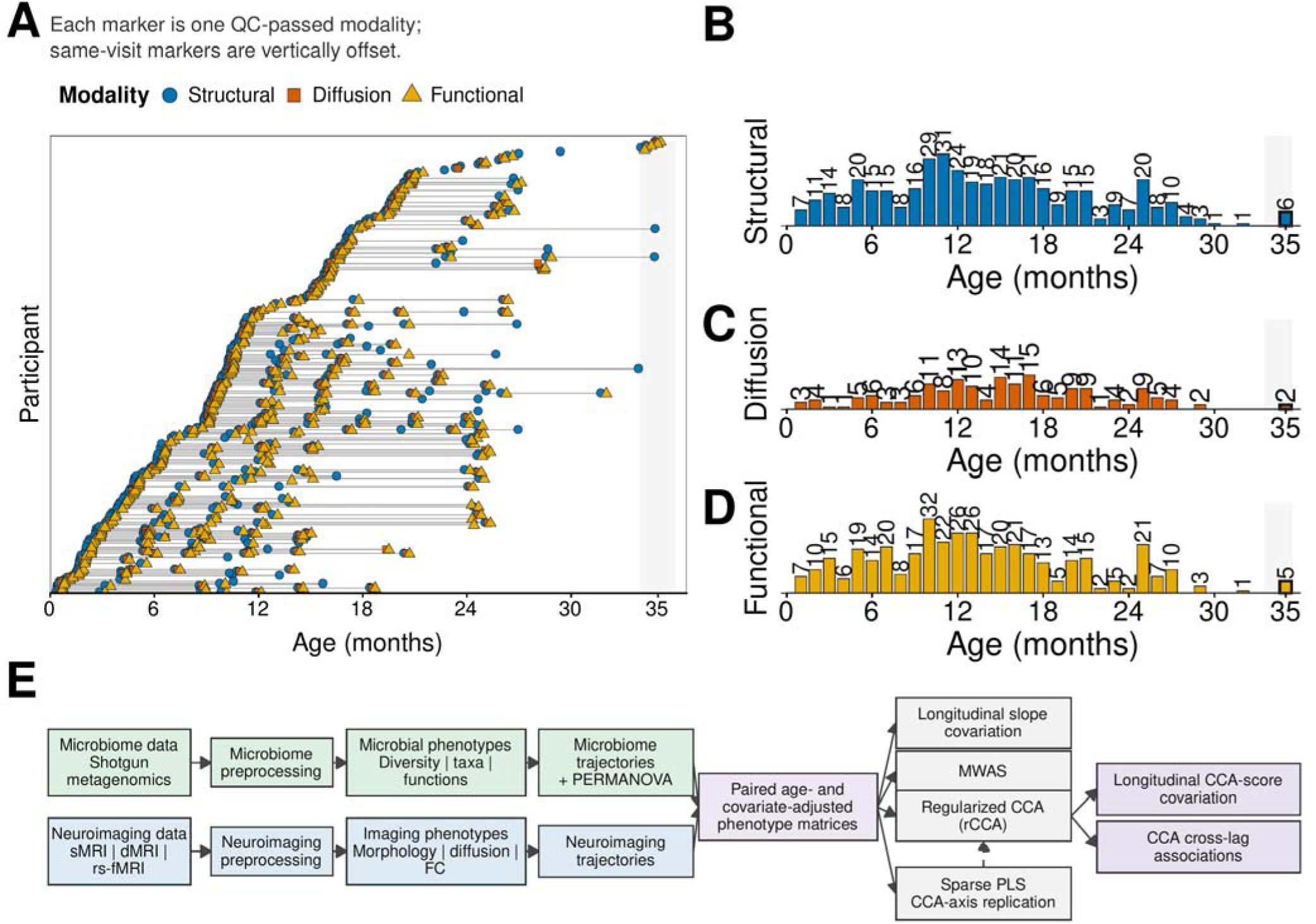
Longitudinal sampling, imaging availability, and analytic workflow. A, QC-passed imaging modalities available at visits with paired fecal samples. Each row represents one participant, lines connect repeated visits, and each colored marker represents one QC-passed modality; markers from the same visit are offset so that all available modalities remain visible. B–D, Monthly counts of QC-passed paired sessions for structural MRI (B), diffusion MRI (C), and resting-state functional MRI (D). Colors match panel A, all age axes begin at 0 months, and the 35-month bin contains 6 structural, 2 diffusion, and 5 functional records, respectively. E, Study workflow. Shotgun metagenomic and structural, diffusion, and resting-state functional MRI data undergo separate preprocessing and phenotype extraction, followed by domain-specific developmental trajectory analyses and microbiome PERMANOVA. The paired age- and covariate-adjusted phenotype matrices are then used for longitudinal slope covariation, microbiome-wide association studies (MWAS), regularized canonical correlation analysis (rCCA), longitudinal CCA-score covariation, CCA cross-lag associations, and sparse partial least-squares (PLS) replication of the CCA axes.

**Table 1.**
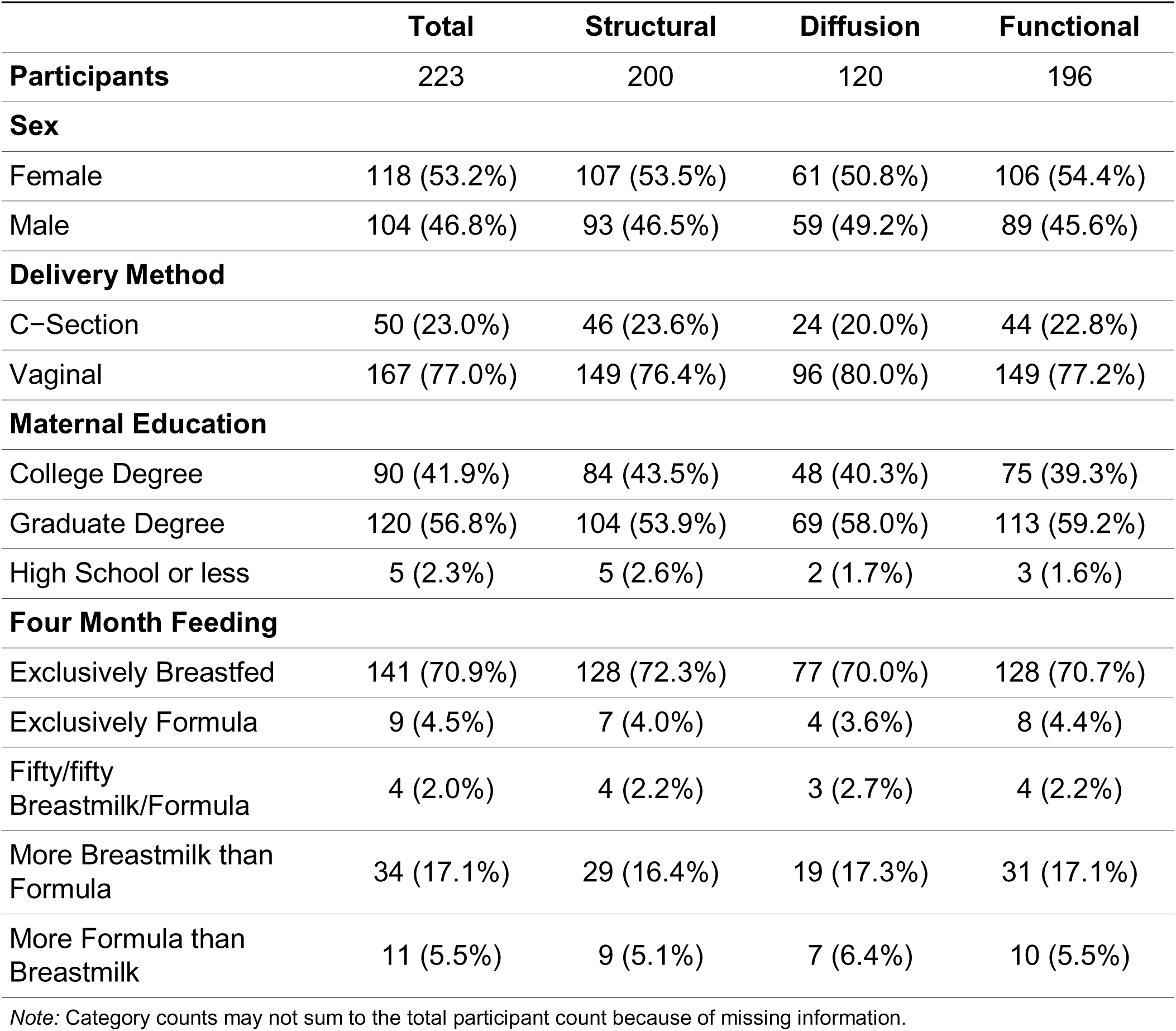
Demographic information of the study cohort.

### Neuroimaging trajectories during early development

We characterized structural, white-matter, and functional development from birth to three years (Figure 2). Most structural measures rose rapidly during the first year, whereas gyrification changed little. FA and MO increased and MD decreased until approximately 20 months, after which changes slowed. Functional networks followed distinct trajectories. Sensorimotor connectivity declined after birth; visual network connectivity increased initially and then declined; and higher-order networks (default mode, salience, executive control) strengthened gradually, with different time courses [12, 13]. These trajectories were used to characterize age-related neuroimaging variation before the cross-domain analyses.

**Figure 2.**
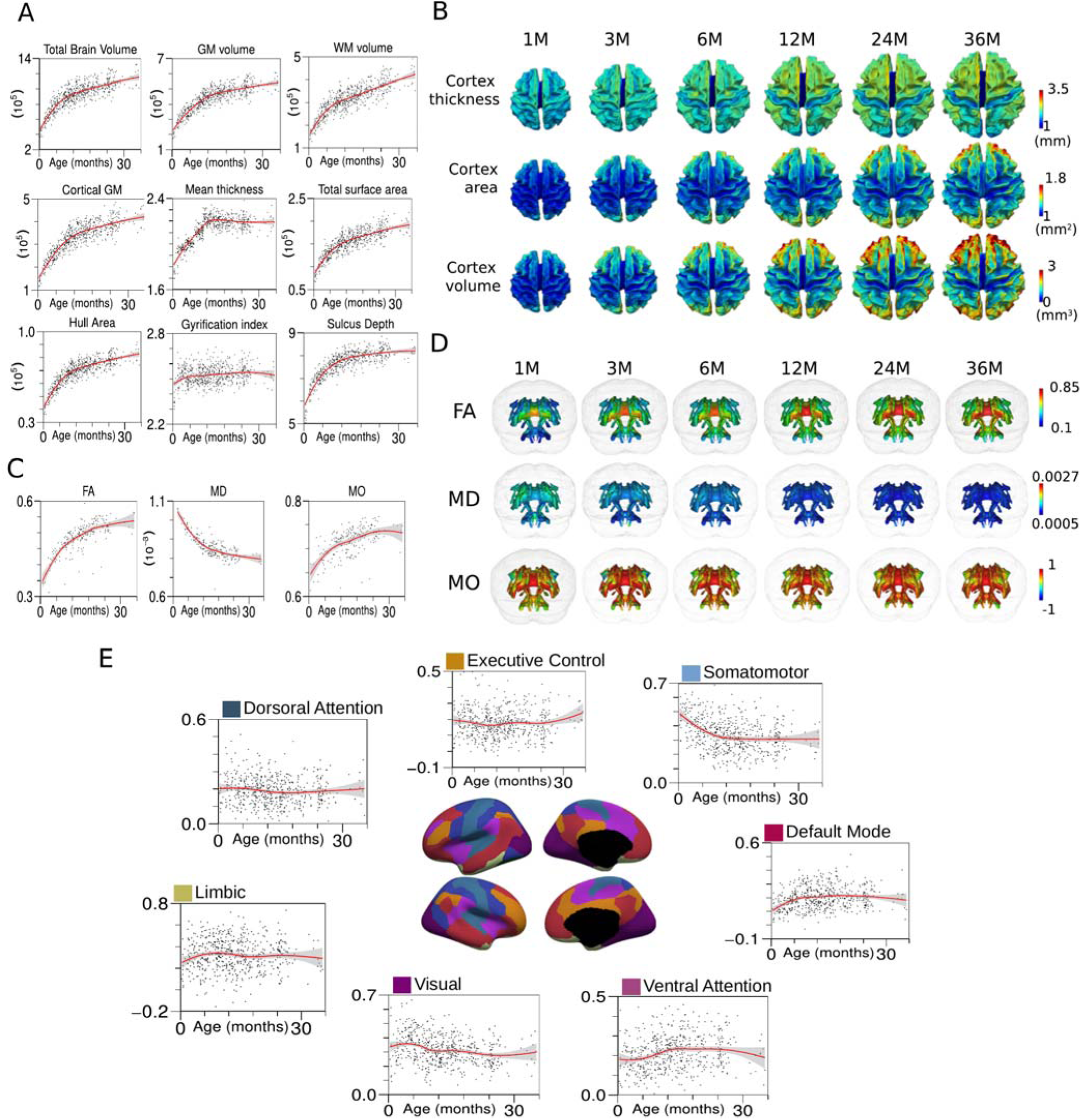
Developmental trajectories of neuroimaging phenotypes from birth to three years. **A**, Whole-brain morphological phenotypes. B, Regional cortical thickness, surface area, and volume at representative ages. C, Mean fractional anisotropy (FA), mean diffusivity (MD), and mode of anisotropy (MO). D, Spatial maps of FA, MD, and MO. E, Seven functional-network trajectories. In trajectory panels A, C, and E, x axes are labeled Age (months) and use rounded limits with unlabeled intermediate ticks. Red curves show fitted developmental trajectories, and gray bands show 95% confidence intervals. Panel E shows mean within-network resting-state functional connectivity rather than somatomotor skill or performance.

### Gut microbiome trajectories during early development

At birth, the predominant phyla were *Actinobacteria*, *Bacteroidetes*, *Firmicutes*, and *Proteobacteria* (Figure 3A), consistent with reported early-life colonization patterns [14]. *Actinobacteria* and *Proteobacteria* declined sharply during the first year as community composition changed with age. This period coincides with reduced breast-milk dependence, dietary diversification, increased environmental exposure, and maturation of the gut environment [15, 16]. After the first year, *Bacteroidetes* and *Firmicutes*—the dominant phyla in healthy adults [17]—increased in relative abundance (Figure 3A) [15, 16]. Across all samples, we observed 684 bacterial species, 57% of which belonged to *Firmicutes* (Supplementary Figure S1). *Bifidobacterium longum* (*B. longum*), *Bifidobacterium breve* (*B. breve*), and *Bifidobacterium bifidum* (*B. bifidum*) were among the most abundant species (Figure 3B and Supplementary Figure S2); bifidobacteria are prominent early-life colonizers [18].

**Figure 3.**
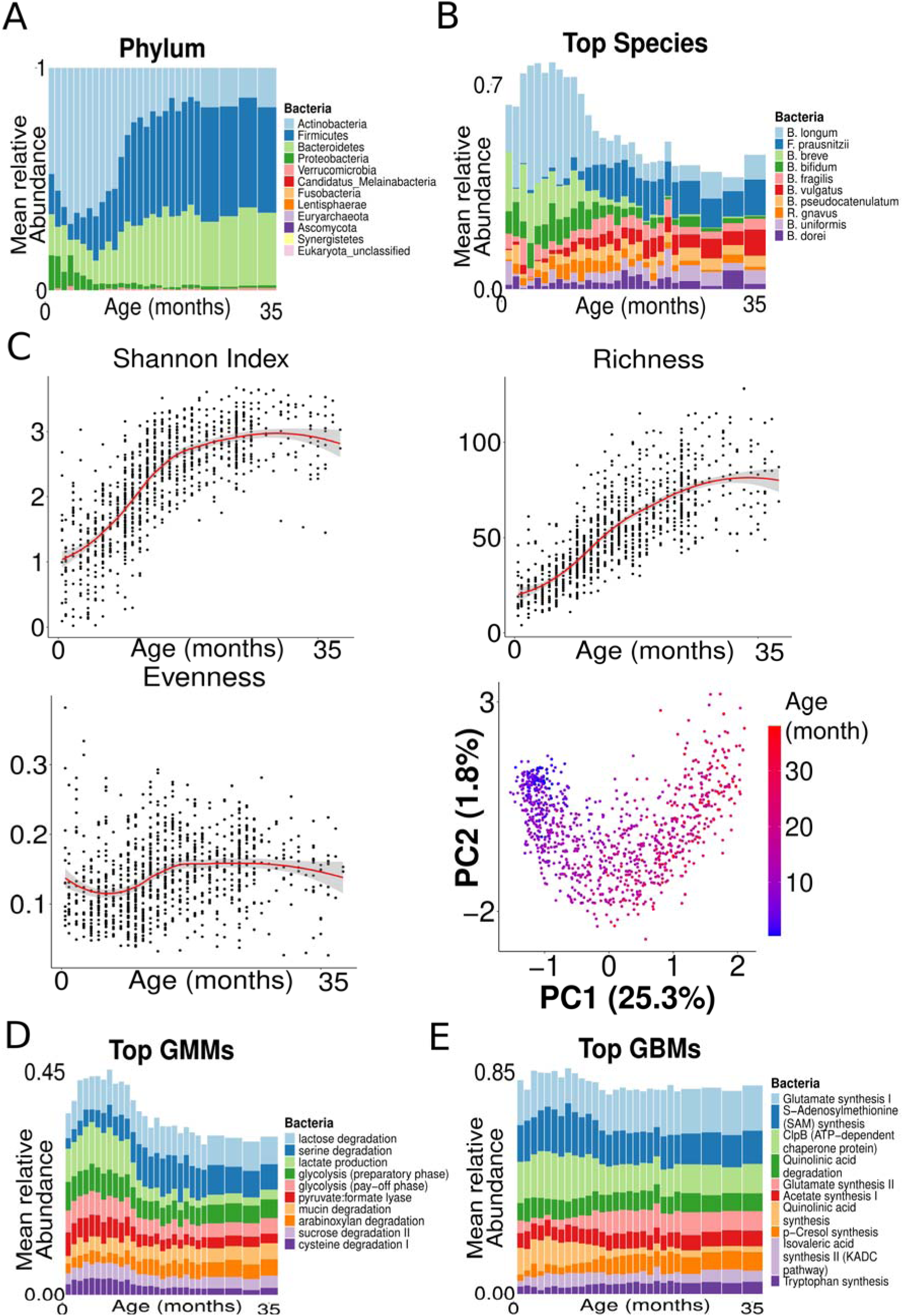
Longitudinal dynamics of gut microbiome composition and diversity during the first three years. A, Mean relative abundance of bacterial phyla, normalized within each age bin. B, Mean relative abundance of the 10 most abundant species; the top 10 are shown for legibility, while Supplementary Figure S2 displays the 100 most abundant species. C, Shannon diversity, richness, evenness, and species-level beta diversity. The beta-diversity panel is an ordination (PC1 versus PC2), not a trajectory plot; point color encodes age in months. D–E, Mean relative abundance of the 10 most abundant gut metabolic modules (GMMs) and gut–brain modules (GBMs). For A, B, D, and E, bars summarize monthly intervals from 0–24 months and three-month intervals thereafter. CLR, centered log-ratio.

Alpha diversity increased with age (Figure 3C): Shannon diversity and richness rose steadily, whereas evenness increased after an initial delay. Species-level ordination also shifted with age while retaining substantial between-child variation. PERMANOVA identified age as the largest measured correlate of beta diversity (R^2^ = 0.107, p < 0.0001). Maternal education, feeding practice at four months, and delivery mode were also associated with beta diversity, each accounting for 1.0–1.5% of variation (p < 0.0001; Methods and Supplementary Text).Functional profiling identified 554 MetaCyc pathways, 44 gut–brain modules (GBMs), and 99 gut metabolic modules (GMMs; Methods). MetaCyc pathways describe microbial reactions, GBMs summarize neuroactive pathways, and GMMs capture core metabolic functions relevant to host nutrition. Figures 3D and 3E show the 10 most abundant GMMs and GBMs; Supplementary Figures S3 and S4 show the complete profiles. GMM abundance declined near 12 months and then plateaued, whereas GBM abundance changed modestly. Lactose degradation and lactate production dominated early GMM profiles before serine degradation became most abundant. Glutamate synthesis, S-adenosylmethionine synthesis, and ClpB remained the three most abundant GBMs across 0–3 years.

### Multilevel microbiome-wide association analysis

We analyzed six microbiome feature sets: five diversity indices, 44 GBMs, 99 GMMs, 202 genera, 684 species, and 554 MetaCyc pathways (Supplementary Table S1). Neuroimaging principal components are summarized in Figure 4. We defined q < 0.05 as statistically significant; associations with 0.05 ≤ q < 0.25 are reported only in the Supplementary Information as exploratory estimates. Five associations met q < 0.05 (Figure 5A). Total brain volume was positively associated with purine deoxyribonucleoside degradation (PWY0-1297; β = 0.22, q < 0.001) and purine nucleotide salvage (PWY66-409; β = 0.15, q = 0.032).

**Figure 4.**
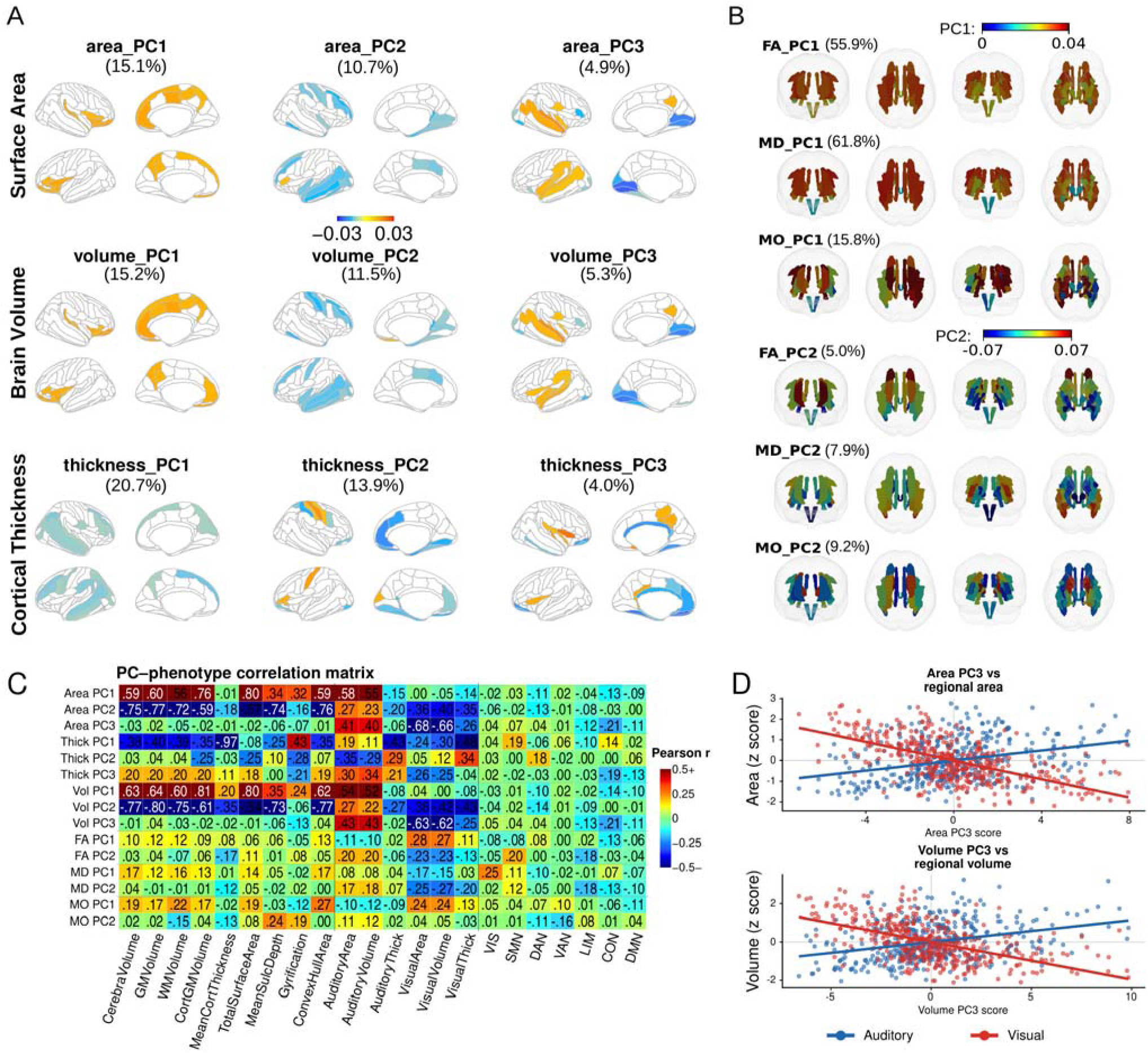
Principal-component structure and phenotypic interpretation of neuroimaging measures. A, Loadings of the top three principal components (PC1–PC3) for cortical surface area, thickness, and volume across 30 cortical regions; percentages indicate variance explained and colors indicate loading coefficients. B, Loadings of the top two components for fractional anisotropy (FA), mean diffusivity (MD), and mode of anisotropy (MO) across white-matter tracts. C, Pearson correlation matrix relating 15 structural and diffusion PCs to nine global cerebral measures, six auditory/visual regional measures, and seven functional-network connectivity measures in 160 matched observations. Values are descriptive correlations from raw measurements without covariate regression; the color scale is capped at r ≤ −0.5 and r ≥ 0.5. D, Observation-level scatterplots of area PC3 with auditory and visual area (top) and volume PC3 with auditory and visual volume (bottom). Regional measures were separately z-standardized for joint display. Blue denotes the auditory region (right transverse temporal sulcus), red denotes the visual region (right occipital pole), and lines are least-squares fits. PC, principal component.

**Figure 5.**
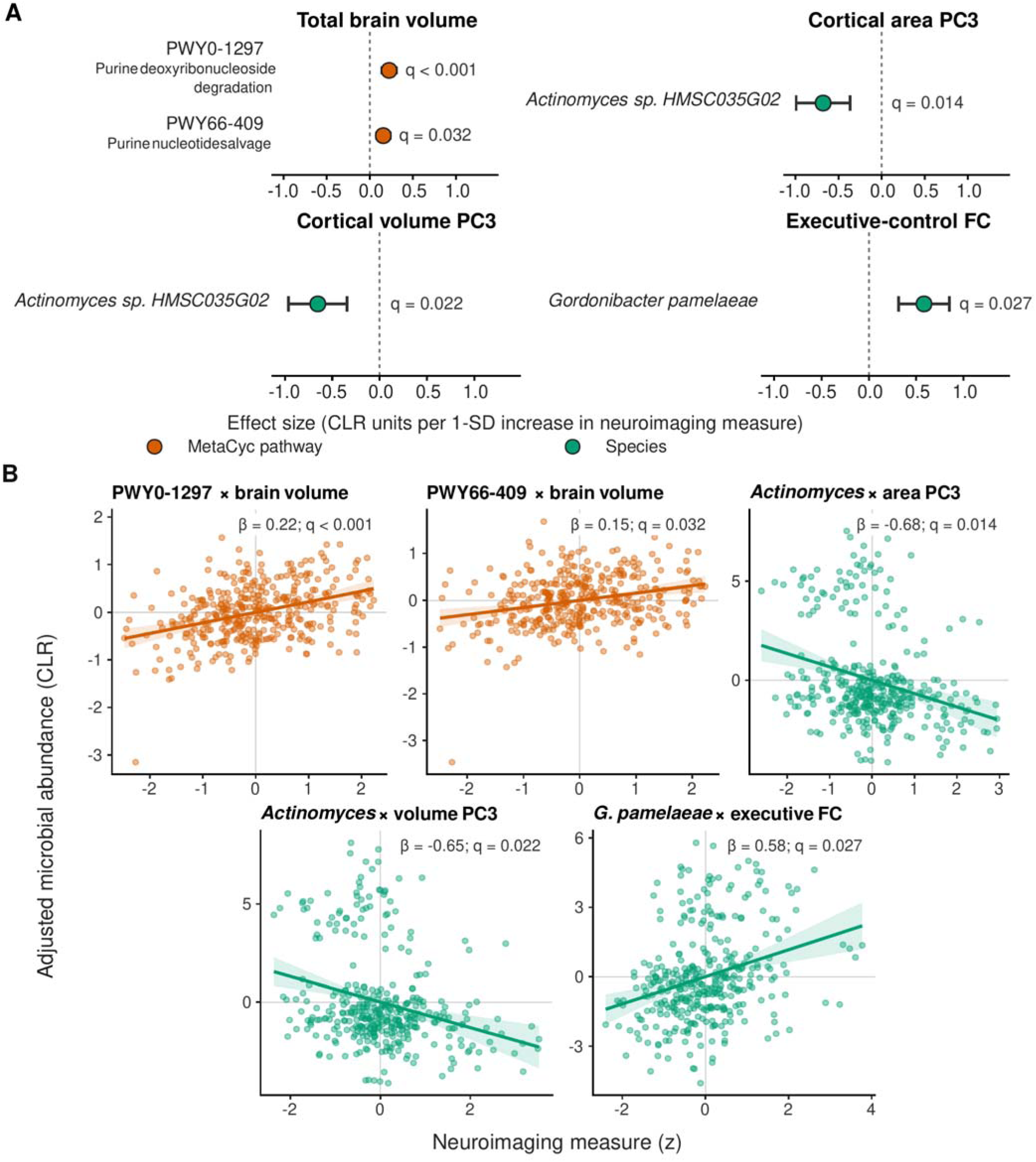
FDR-significant associations between neuroimaging measures and gut microbial features. A, Effect estimates and 95% confidence intervals for all five associations with q < 0.05. Effect sizes are CLR units per 1-SD increase in the neuroimaging measure; the dashed vertical line denotes no association. Orange denotes MetaCyc pathways and green denotes species. B, Covariate-adjusted partial-residual plots for the same five associations. Points represent matched stool–MRI observations, solid lines show the focal fitted associations, and shaded bands show % confidence intervals. The β estimates and q values correspond to the models in A. Continuous neuroimaging predictors were z-standardized, and microbial abundances were CLR-transformed. FDR, false discovery rate; CLR, centered log-ratio; PC, principal component; FC, functional connectivity.

*Actinomyces sp. HMSC035G02* abundance was inversely associated with cortical area PC3 (β = −0.68, q = 0.014) and cortical volume PC3 (β = −0.65, q = 0.022), whereas *Gordonibacter pamelaeae* abundance was positively associated with executive-control connectivity (β = 0.58, q = 0.027). Effect sizes are CLR units per 1-SD increase in the neuroimaging measure.

Covariate-adjusted partial-residual plots show the direction and observation-level dispersion of these five associations (Figure 5B). Supplementary Excel Table S1 provides the complete MWAS statistics.

### Covariation of microbiome and cortical growth rates

We tested whether within-participant developmental rates covaried by estimating covariate-adjusted age slopes for each microbial and neuroimaging feature. We excluded a participant from a slope pair when either slope lay at least five median absolute deviations from its median. Two pairs met the prespecified within-family FDR threshold (Figure 6). The allose-degradation GMM slope correlated positively with cortical-thickness PC1 (n = 102, r = 0.47, 95% CI 0.31–0.61, q = 0.000424), whereas the tyrosine-degradation-II GMM slope correlated inversely with surface-area PC2 (n = 102, r = −0.37, 95% CI −0.53 to −0.19, q = 0.0417).

**Figure 6.**
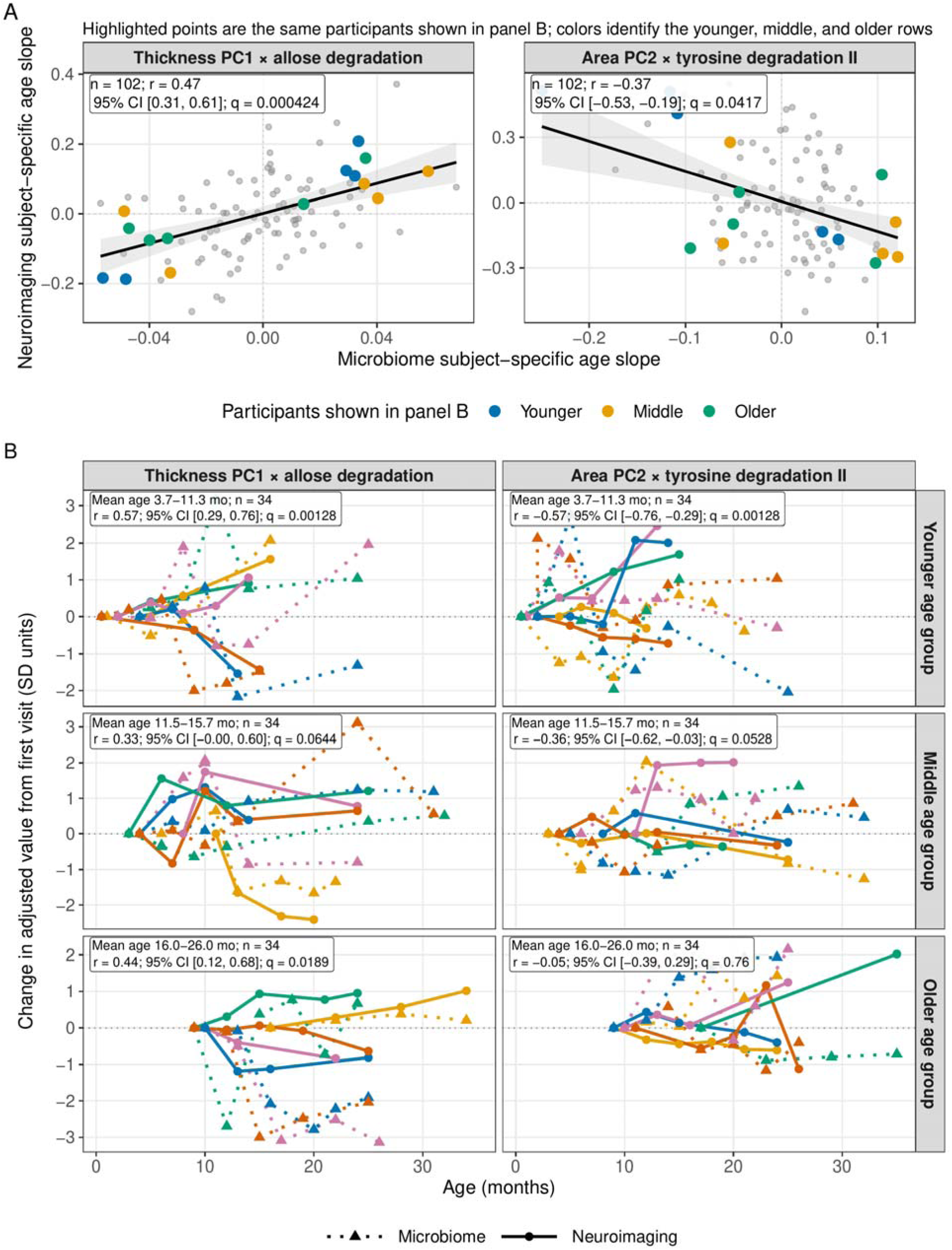
Covariation of participant-specific microbiome and cortical growth rates. A, Pearson correlations between covariate-adjusted participant-specific age slopes. Gray points show all participants retained after the five-MAD filter; colored points identify the same participants shown in B (blue, younger age group; orange, middle age group; green, older age group). Black lines show linear fits and shaded bands show 95% confidence intervals. B, Longitudinal profiles for five representative participants per mean-age group, selected for long follow-up, marked growth or decline, and proximity to the fitted slope relation. Dotted lines denote the microbial feature and solid lines denote the neuroimaging feature, with features from the same participant shown in the same color. Text boxes report correlations calculated from all retained participants in each age group, not only those displayed; age-group q values are FDR-adjusted across the six displayed tests. Microbial abundance features were CLR-transformed. Participant-specific slopes were adjusted for the prespecified covariates, and total brain volume was additionally included for the surface-area outcome. PC directions should be interpreted from the regional loadings in Figure 4. MAD, median absolute deviation; GMM, gut metabolic module; FDR, false discovery rate.

The allose-degradation association was positive in all three mean-age groups, although the middle group did not meet the FDR threshold for the displayed age-stratified tests. The tyrosine-degradation association was strongest in the younger group, remained negative in the middle group, and approached zero in the older group. These estimates are not consistent with a constant association across infancy. PC scores reflect regional loading patterns rather than global anatomical change; for example, a higher cortical-thickness PC1 score represents stronger expression of its negative-loading pattern rather than generalized cortical thickening (Figure 4).

### Gut–brain covariance axes for brain structure and function

We applied ridge-regularized canonical correlation analysis to 107 children with complete data for 78 microbial and 22 neuroimaging measures (Methods; Supplementary Tables S4 and S11). Ten-fold cross-validation selected the ridge penalties. Significance was evaluated with 1,000 permutations, and generalization was assessed in 1,000 random 75%/25% train/test splits. CCA1 and CCA2 exceeded the permutation envelope and yielded positive held-out correlations in 95% and 86% of splits, respectively. Only these two modes were carried forward for interpretation (Figure 7).

**Figure 7.**
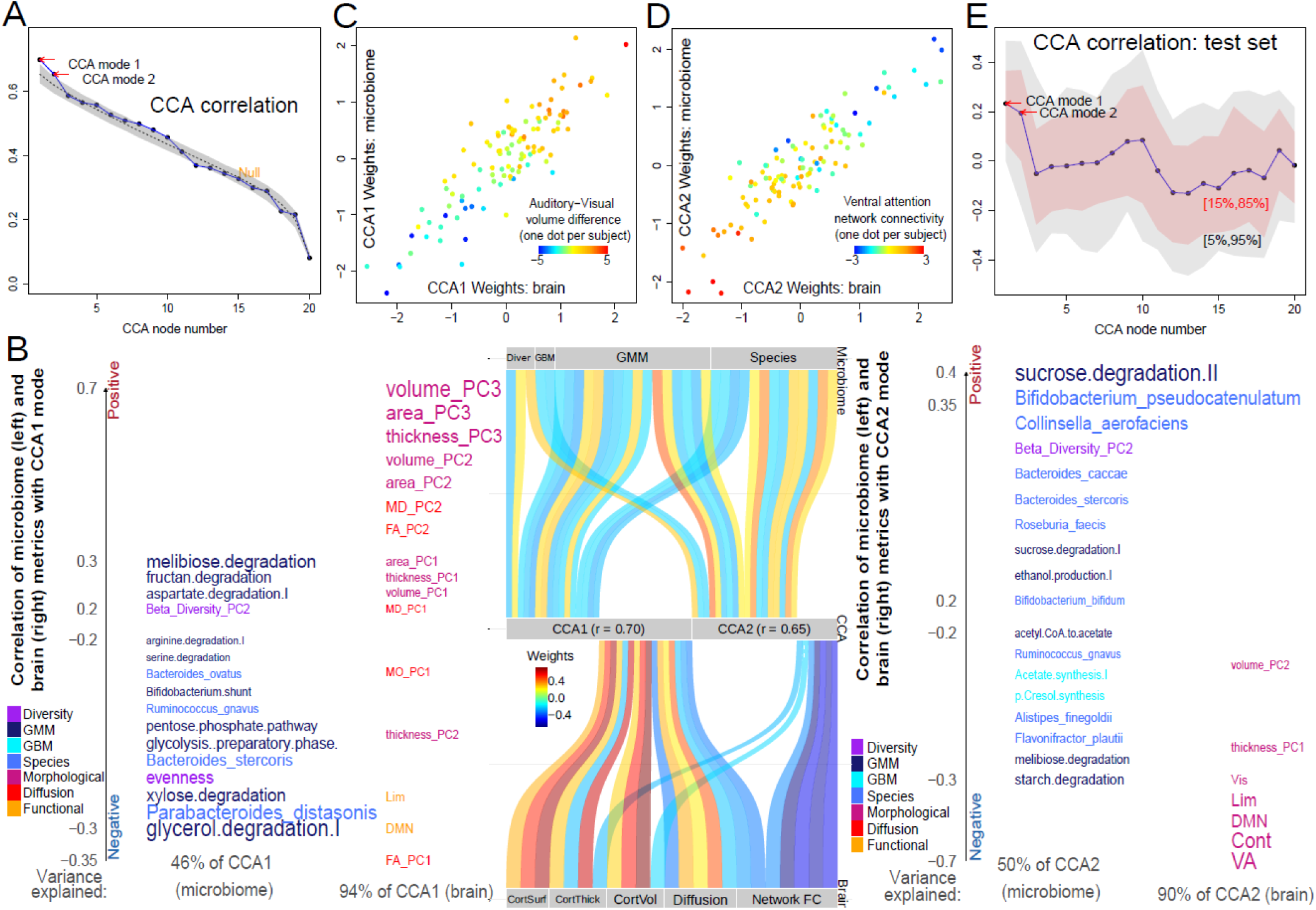
Regularized canonical correlation analysis (rCCA) of gut microbiome and neuroimaging measures. **A**, The first 20 canonical pairs are shown in descending order of canonical correlation; 20 components were displayed for visualization and did not define the inferential cutoff. Modes were retained only when they exceeded the 95th percentile of the permutation null and showed a positive held-out correlation in more than 85% of 1,000 Monte Carlo splits. B, Variable contributions to CCA1 and CCA2. C–D, Participant-level canonical scores for CCA1 and CCA2. E, Held-out correlations across 1,000 random 75%/25% train/test splits; ribbons show empirical intervals. Only the first two modes met the prespecified stability criterion.

CCA1 primarily linked structural measures—volume PC3 (r = 0.71), area PC3 (r = 0.66), and thickness PC3 (r = 0.60)—with microbial functions that included melibiose and fructan degradation. The raw-measurement correlations in Figure 4C–D clarify the direction of the structural PC3 scores: area PC3 correlated positively with auditory area (r = 0.41) and negatively with visual area (r = −0.68), whereas volume PC3 correlated positively with auditory volume (r = 0.43) and negatively with visual volume (r = −0.62). Higher area or volume PC3 scores therefore index a relative auditory–visual regional contrast rather than a global increase in cortical size. We refer to this pattern as the structural–metabolic axis. The label reflects the dominant loadings rather than an exclusive mapping: taxonomic features also contributed, and CCA does not establish a causal effect of microbial metabolism on brain structure. Cortical area PC3 and volume PC3 were also the structural outcomes associated with *Actinomyces sp. HMSC035G02* in the FDR-significant MWAS (Figure 5). Because both PCs encode the same auditory–visual regional contrast (Figure 4C–D), the feature-level and multivariate analyses point to the same structural phenotype within this cohort. This agreement supports the anatomical interpretation of CCA1 but is not independent validation. *Actinomyces* species are facultative anaerobic commensals of the oral cavity, gastrointestinal tract, and genitourinary tract; under pathological conditions, some species can act as opportunistic pathogens and have been associated with systemic inflammatory responses.

CCA2 primarily linked species composition and GBMs—including *Bifidobacterium pseudocatenulatum* (r = 0.37) and sucrose degradation II (r = 0.42)—with functional connectivity, most strongly in the ventral-attention network (r = −0.68). In the displayed orientation, higher microbial CCA2 scores accompanied weaker ventral-attention, executive-control, and default-mode connectivity. Because canonical signs are arbitrary, this orientation does not imply that the contributing taxa reduce connectivity. We refer to this pattern as the functional–taxonomic axis. The axis name summarizes the strongest loadings and does not imply biologically exclusive feature sets.

The rCCA and feature-level growth analyses capture different aspects of cross-domain covariation. rCCA summarizes participant-level covariance across features, whereas the growth analysis compares within-participant rates of change. Both significant slope pairs linked microbial metabolic modules to cortical morphology, placing them in the same broad feature classes as CCA1. The specific pathway–component pairs differed, however, and do not validate CCA1, its PC3 sensory contrast, or a mechanism. We therefore examined persistence and temporal ordering separately using longitudinal canonical scores.

### Longitudinal covariation and cross-lag associations of the CCA axes

We projected eligible longitudinal observations onto the frozen CCA weights and estimated adjusted age slopes for each domain. Among 29 children with repeated measures in both domains, neither matched slope correlation survived FDR correction (CCA1: r = 0.04, 95% CI −0.33 to 0.40, q = 0.825; CCA2: r = 0.25, 95% CI −0.13 to 0.56, q = 0.390; Supplementary Figure S13). Age-stratified correlations were also nonsignificant (all q ≥ 0.269). The data therefore did not show significant covariance between rates of change in microbiome and neuroimaging canonical scores.

Autoregressive cross-lag models tested both directions across 1–3-, 3–6-, and 6–12-month windows. Earlier neuroimaging CCA1 predicted later microbiome CCA1 over 1–3 months (standardized β = 0.35, 95% CI 0.11–0.59, p = 0.0054, q = 0.0327; 65 pairs from 40 children) and 3–6 months (β = 0.31, 95% CI 0.06–0.56, p = 0.0148, q = 0.0444; 92 pairs from 59 children). No reverse-direction CCA1 association or CCA2 association survived FDR correction (Figure 8). This pattern is consistent with temporal precedence within the measured windows but does not establish causality.

**Figure 8.**
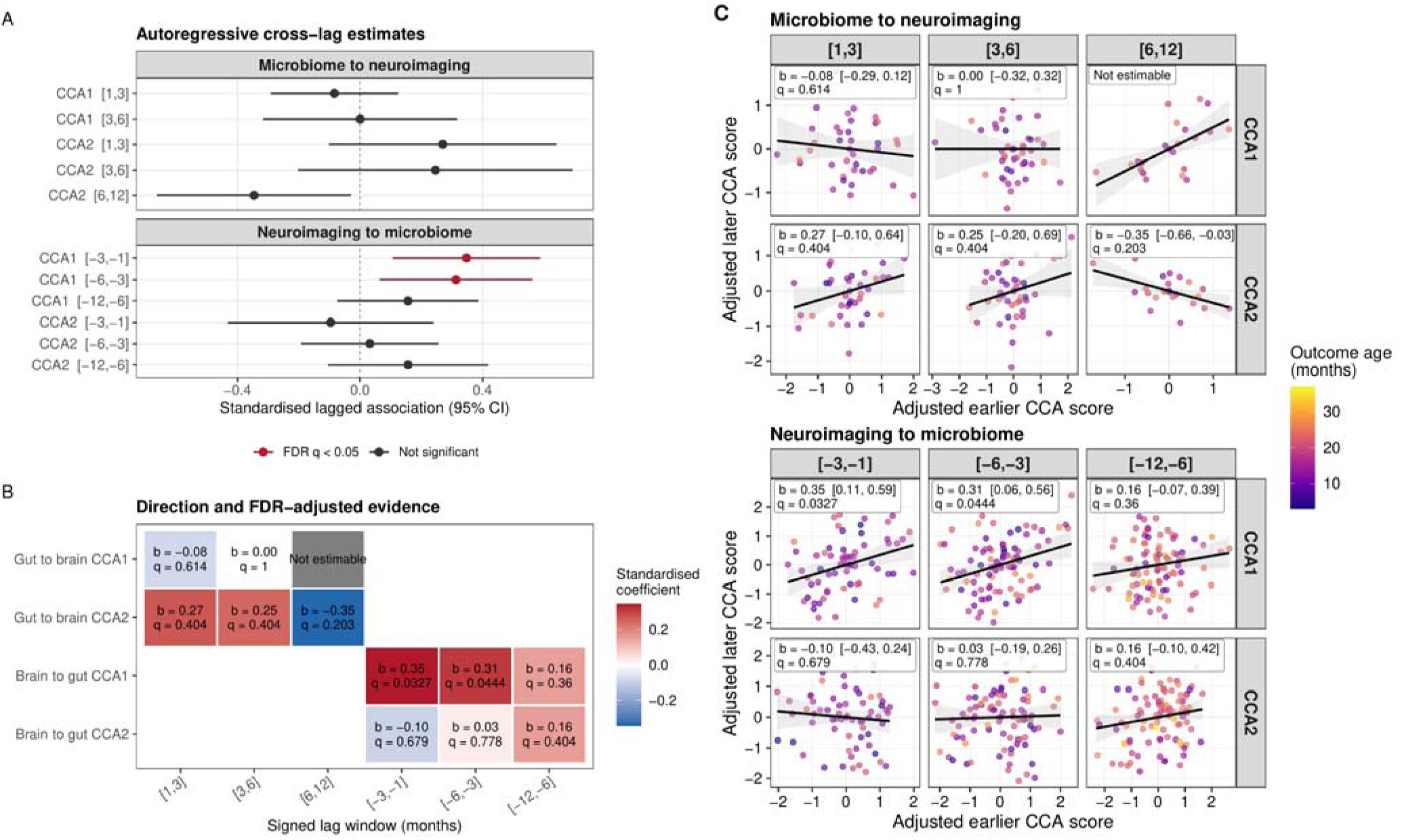
Bidirectional cross-lag associations of longitudinal CCA scores. A, Standardized autoregressive cross-lag estimates and 95% confidence intervals for CCA1 and CCA2 across prespecified lag windows. Red denotes FDR q < 0.05. B, Heat map of effect estimates and FDR-adjusted q values; positive signed lags indicate earlier microbiome and later neuroimaging observations, whereas negative signed lags indicate earlier neuroimaging and later microbiome observations. C, Covariate-adjusted earlier and later CCA scores for each direction, mode, and lag window; lines and shaded bands are descriptive linear fits with 95% confidence intervals, and point color denotes outcome age. Models adjusted for the most recent prior outcome score, response age and age squared, sex and age-by-sex terms, site, maternal education, delivery mode, feeding practice at four months, sequencing depth, total brain volume, and exact lag duration. FDR correction was applied within each canonical mode across the six directional lag-window tests.

### Cross-method evaluation of CCA axes with sparse partial least squares

To test whether the CCA structure was recoverable with a distinct multivariate method, we fitted sparse canonical partial least squares (PLS) to the same 107 participants and the same residualized 78-variable microbiome and 22-variable neuroimaging blocks (Methods; Figure 9). Across 20 native PLS components, joint correspondence across the two blocks was strongest for CCA1 with PLS1 (mean score correlation r = 0.55) and PLS2 (r = −0.52), and for CCA2 with PLS5 (r = 0.70; Figure 9A–C). Examined separately, PLS1 and PLS2 correlated with CCA1 scores in the microbiome (r = 0.37 and −0.56) and neuroimaging (r = 0.74 and −0.48) blocks, whereas PLS5 correlated with CCA2 scores in the microbiome (r = 0.55) and neuroimaging (r = 0.85) blocks. The negative PLS2 correlations reflect the arbitrary orientation of latent-variable signs rather than a discordant biological direction.

**Figure 9.**
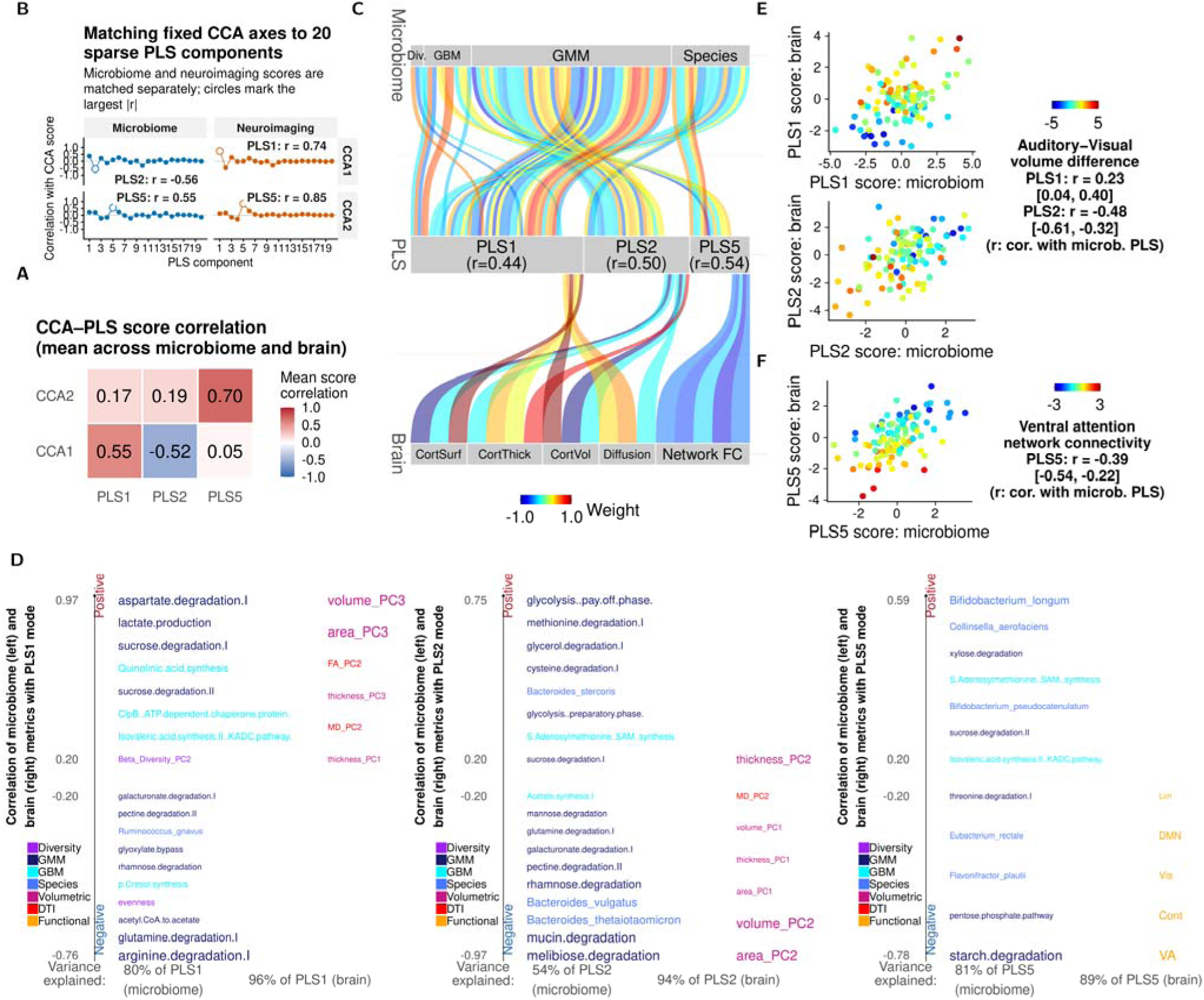
Sparse canonical partial least-squares replication of the CCA axes. A, Mean correlations between fixed CCA scores and the selected native PLS scores, averaged across the microbiome and neuroimaging blocks. B, Correlations between each fixed CCA score and 20 native PLS components, evaluated separately by block; open circles mark the largest absolute correlation in each facet. C, Alluvial summary of microbiome and neuroimaging feature classes contributing to PLS1, PLS2, and PLS5; link color denotes the signed feature–score correlation, and values in the central strips are the microbiome–neuroimaging PLS score correlations. D, Microbiome (left) and neuroimaging (right) features correlated with PLS1, PLS2, and PLS5; vertical position gives direction and magnitude, text color indicates feature class, and labels are shown for correlations with |r| ≥ 0.20 (or the strongest features when fewer met that threshold). E, Participant-level microbiome and neuroimaging scores for PLS1 and PLS2, colored by auditory–visual volume difference; annotations report correlations between the phenotype and microbiome PLS scores with 95% confidence intervals. F, Corresponding PLS5 scores colored by ventral-attention network connectivity. Latent-variable signs are arbitrary; PLS2 has the inverse orientation to CCA1.

The same phenotype-level patterns were evident in the PLS solution. Microbiome PLS1 scores correlated positively with the auditory–visual volume difference (r = 0.23, 95% CI 0.04–0.40, p = 0.0167), whereas PLS2 scores correlated inversely with this contrast (r = −0.48, 95% CI −0.61 to −0.32, p = 1.88 × 10□□). Microbiome PLS5 scores correlated inversely with ventral-attention network connectivity (r = −0.39, 95% CI −0.54 to −0.22, p = 2.55 × 10□□; Figure 9E–F). The PLS solution therefore reproduced the structural contrast defining CCA1 and the functional-network pattern defining CCA2.

Because PLS used the same participants and residualized variables, it provides within-sample cross-method corroboration rather than external validation. In 1,000 repeated participant-level 90%/10% training–test splits (96 training and 11 test participants), each component refit was matched one-to-one to full-data reference PLS1, PLS2, and PLS5 by maximizing the joint absolute cosine similarity of the microbiome and neuroimaging loading vectors. Mean training cross-block correlations were 0.472, 0.504, and 0.562, and mean held-out test correlations were 0.063, 0.144, and 0.091 for PLS1, PLS2, and PLS5, respectively. Held-out correlations were positive in 59%, 68%, and 64% of the 1,000 splits, respectively. The full-sample PLS modes captured the dominant CCA score and feature patterns but showed limited split-sample generalization.

## Discussion

During the first three years of life, brain phenotypes and gut microbial composition changed substantially with age. Cross-sectional multivariate analyses identified structural–metabolic and functional–taxonomic covariance patterns, while feature-level longitudinal analyses identified two metabolic–cortical slope associations. Canonical-score slopes were not significantly correlated. In contrast, neuroimaging CCA1 predicted microbiome CCA1 1–6 months later in cross-lag models. The findings indicate coordinated variation across domains and temporal ordering for CCA1 but do not establish causal direction.

At the prespecified q < 0.05 threshold, two purine-recycling pathways were positively associated with total brain volume, *Actinomyces sp. HMSC035G02* was inversely associated with cortical area and volume PC3 scores, and *Gordonibacter pamelaeae* was positively associated with executive-control connectivity. Figure 5B displays the adjusted observation-level distributions for all five findings; the broader q < 0.25 results remain in the Supplementary Information as hypothesis-generating evidence.

The *Gordonibacter pamelaeae* association with executive-control connectivity remains mechanistically unresolved. Cultured *G. pamelaeae* can convert ellagic acid to intermediate urolithins [19], making diet-derived urolithin metabolism a plausible target for follow-up. We did not measure substrate availability, metabolite concentrations, or mediation. Infant metabolomics and prospective sampling will be required before culture-based or adult findings can be extended to this cohort.

The inverse associations of *Actinomyces sp. HMSC035G02* with cortical area PC3 and volume PC3 should not be interpreted as reduced global cortical growth because each PC3 score represents a multiregional contrast rather than total cortical size. Their biological basis remains unknown. The positive associations of purine salvage and deoxyribonucleoside degradation with total brain volume warrant direct measurement of purine metabolites in future studies. Species abundance does not establish organismal activity, and pathway abundance reflects genetic potential rather than metabolite flux; mechanistic interpretation is therefore premature. The same two PC3 outcomes associated with *Actinomyces sp. HMSC035G02* were also among the strongest structural contributors to CCA1 and encode the auditory–visual contrast in Figure 4C–D. The feature-level MWAS and multivariate CCA therefore identify the same sensory-regional structural phenotype within this cohort. This agreement supports the CCA1 interpretation but does not constitute independent replication.

The structural–metabolic pattern links cortical morphology to contrasting microbial carbohydrate functions. Surface-area and volume PC3 loadings oppose lateral temporal and perisylvian regions to medial occipital regions, producing the auditory–visual contrast shown in Figures 4A and 7B–C. Thickness PC3 is more spatially heterogeneous, and other structural and diffusion components also contribute. The anatomical interpretation therefore centers on the temporal–occipital contrast rather than uniform morphological change. Microbial loadings likewise contrast melibiose and fructan degradation with glycolysis preparation, the pentose phosphate pathway, and the Bifidobacterium shunt. They indicate contrasting metabolic potential, not uniformly increased glycolysis or short-chain-fatty-acid production.

Fermentation products are one plausible route by which microbial metabolism could relate to cortical development. In mice, microbial short-chain fatty acids regulate microglial maturation [20], and microbiota manipulation or short-chain-fatty-acid administration can alter adult visual-cortical plasticity [21]. These animal findings suggest a possible metabolite–glia–plasticity pathway but do not account for the MRI pattern observed here. Metabolic or immune signals may also relate differently to cortical regions with distinct developmental schedules [22]. Participant-level PC scores cannot establish accelerated maturation, selective microbial effects on sensory cortex, or the full spatial pattern of CCA1.

The functional–taxonomic pattern links microbial composition to distributed network connectivity. *Bifidobacterium pseudocatenulatum*, *Collinsella aerofaciens*, and sucrose- and acetate-related modules contributed alongside ventral-attention, executive-control, default-mode, limbic, and visual connectivity. Their opposing loadings do not show that either species reduces connectivity. Taxonomic variation may instead reflect substrate use and cross-feeding that alter community output. Coculture studies show cross-feeding between *Bifidobacterium adolescentis* and butyrate producers [23], and mouse studies show vagus-dependent microbial modulation of central GABA-receptor expression [24]. These mechanisms remain untested here. The axis names therefore describe statistical contributions, not separate biological systems.

Sparse PLS provided a methodologically distinct check on the CCA patterns, with signs aligned jointly across the microbiome and neuroimaging blocks rather than defined by either block alone. At |r| ≥ 0.20, fructan degradation and *Bacteroides stercoris* had the same direction in CCA1, PLS1, and sign-aligned PLS2. CCA1–PLS1 concordance additionally included aspartate degradation I, xylose degradation, evenness, beta-diversity PC2, *Ruminococcus gnavus*, and several other microbial features; CCA1–PLS2 concordance included glycerol degradation I, melibiose degradation, and the preparatory phase of glycolysis. Melibiose degradation was the only shared CCA1 feature with the opposite direction in PLS1, indicating substantial but incomplete agreement. For CCA2 and PLS5, all five shared microbiome features above this threshold were directionally concordant: *Collinsella aerofaciens*,

*Bifidobacterium pseudocatenulatum*, and sucrose degradation II were positive, whereas starch degradation and *Flavonifractor plautii* were negative. The corresponding brain patterns were also concordant: PLS1 emphasized the positive CCA1 volume-PC3 and area-PC3 contrast, sign-aligned PLS2 emphasized positive area-PC2 and volume-PC2 contributions, and PLS5 reproduced the negative CCA2 ventral-attention, executive-control, visual, default-mode, and limbic connectivity contributions. Agreement across both data blocks makes it less likely that either CCA axis reflects a pattern confined to a single block. However, because PLS used the same participants and residualized variables, it cannot be considered independent replication; external multimodal infant cohorts remain necessary.

The feature-level growth results align most closely with CCA1 because both significant pairs linked microbial metabolic potential to cortical morphology. This alignment concerns feature classes, not the same pathway or component combinations: the longitudinal pairs involved allose degradation, tyrosine degradation II, thickness PC1, and area PC2 rather than CCA1’s defining carbohydrate pathways and PC3 contrast. No functional-connectivity growth association met q < 0.05. The feature-level results therefore support metabolic–cortical covariation but do not establish temporal order, which we tested separately with canonical scores.

Longitudinal canonical scores provided a separate test of the CCA1 pattern over time. Slopes did not show uniform contemporaneous coupling, but higher neuroimaging CCA1 predicted higher microbiome CCA1 1–3 and 3–6 months later after adjustment for prior microbiome CCA1 and measured covariates. Age-dependent feeding, behavior, gastrointestinal physiology, or other unmeasured exposures could account for this temporal pattern. No reverse-direction CCA1 or CCA2 association survived FDR correction, so the data did not support a general bidirectional lagged pattern. Because canonical signs are arbitrary and the design is observational, these estimates indicate prediction in the fixed score orientation rather than a causal brain-to-microbiome effect.

Two prior longitudinal studies are relevant to these findings. Portlock et al. [25] integrated fecal microbiota and plasma lipids with brain activity and cognition in children with malnutrition.

Bonham et al. [26] linked microbial genes involved in neuroactive-compound metabolism to concurrent and later visual-evoked responses during the first 18 months. Both studies motivate further study of microbial metabolism and sensory development, but they differ from the present MRI rCCA, growth, and cross-lag analyses in population, age range, neural phenotype, and analytic unit. Neither therefore constitutes direct replication.

Several limitations should be considered. The cohort comprised healthy children with limited demographic heterogeneity, and q values from 0.05 to 0.25 were treated as exploratory. The observational design does not establish causality. Cross-lag models provide temporal information within prespecified windows, but irregular sampling and residual confounding remain possible. MRI and stool observations were paired within a maximum of 30 days.

Sensitivity MWAS using 20- and 15-day maximum windows and adjustment for the signed stool-minus-MRI age difference showed broadly stable effect sizes for the five original MWAS signals, but fewer associations retained FDR significance as analytic samples decreased (Supplementary Figure S14 and Supplementary Table S11). The narrower windows therefore reduced precision and power. Available data lacked time-varying breastfeeding status, recent illness, gastrointestinal symptoms, antibiotic exposure, detailed dietary patterns, and a head-motion variable for the longitudinal CCA models; their omission may leave residual confounding. Future longitudinal studies should measure these exposures repeatedly alongside closely synchronized stool and MRI assessments so that they can be modeled as time-varying confounders or moderators. Natural-sleep functional connectivity may also vary with sleep state, and metagenomic profiles measure encoded potential rather than pathway activity or host exposure. Further work should combine targeted metabolomics, independent external replication, additional resampling-based validation, quasi-experimental mediation, and experimental models before directional mechanisms are inferred.

## Methods

### Ethics

The Institutional Review Boards of the University of North Carolina at Chapel Hill and the University of Minnesota approved the study, and parents provided written informed consent. The Supplementary Text details MRI acquisition and participant preparation. Briefly, we scanned infants without sedation during natural sleep on Siemens 3 T Prisma systems with a 32-channel coil and acquired structural T1w/T2w, diffusion, and T2*-weighted resting-state fMRI. Here, ‘resting state’ denotes spontaneous BOLD acquisition during monitored natural sleep rather than wakeful task-free rest.

### DNA sequence quality control

All DNA extraction, library preparation and sequencing were performed by CosmosID, Inc. (Germantown, MD, USA; now Cmbio), a CLIA-certified, ICH-GCP-compliant laboratory (Supplementary Information). We adapted previously reported quality-control procedures based on the Human Microbiome Project protocol [27, 28]. We removed duplicate paired-end reads with cd-hit-dup (-u 50) [29, 30], then used BBDuk to remove Nextera XT adapters and PhiX sequences (ftm=5 tpe tbo qtrim=rl trimq=25 minlen=50 ref=adapters,phix). BBSplit removed reads mapping to GRCh38.p14, and BBDuk filtered low-complexity reads (entropy=0.01 entropywindow=50 entropyk=5). Post-QC depth ranged from 3 to 118 million reads (mean, 17.9 million). We excluded 16 samples with fewer than 3 million reads.

### Taxonomic and functional profiling

We used MetaPhlAn 3.0.2 and HUMAnN for taxonomic and functional profiling, respectively [31]. HUMAnN identified 8,560 UniRef90 gene families [32], which we regrouped into 7,123 KEGG Orthology groups and 554 MetaCyc pathways [33]. We retained pathways and KO groups present in at least 90% of samples. The omixer-rpm R package converted KO profiles into 44 gut–brain modules and 99 gut metabolic modules [34–36]. These module frameworks summarize microbial neuroactive and core metabolic functions.

### Summary-level microbiome features

We calculated Shannon diversity, Simpson evenness, and Chao1 richness from species relative abundances with the vegan R package. To characterize beta diversity, we centered-log-ratio transformed MetaPhlAn species counts and applied principal component analysis with the default prcomp settings. We plotted PC1 and PC2 against sample age; because the PCs are computed on CLR-transformed compositions, Euclidean distances in the PC1/PC2 plane approximate the Aitchison distances between compositions [37].

### Structural neuroimaging processing

Each included structural scan had T1w and T2w images, which we processed with iBEAT V2.0 [38]. We corrected intensity inhomogeneity with N3 [39], aligned T2w to T1w images with FLIRT [40], and removed noncerebral tissue with a convolutional neural network [41]. A densely connected U-Net segmented white matter, gray matter, and cerebrospinal fluid [41, 42]. We separated the hemispheres with an age-matched template, filled subcortical regions and ventricles, corrected topology, and reconstructed inner and pial surfaces [43–46].

Spherical registration aligned each surface to an infant atlas, after which we propagated the 148-region FreeSurfer Destrieux parcellation [47–49]. Visual inspection confirmed cortical folds and parcel boundaries.

For each scan, we calculated total brain volume and nine global cerebral measures: cerebrum, gray-matter, white-matter, and cortical gray-matter volumes; mean cortical thickness; total cortical surface area; mean sulcal depth; mean gyrification index; and convex-hull area. We also measured mean cortical thickness, surface area, and cortical volume within each of the 148 Destrieux regions.

### Functional MRI processing

We preprocessed resting-state fMRI with FSL. We removed the first 10 volumes, corrected motion, and applied 0.01–0.08-Hz band-pass filtering. Boundary-based registration aligned each series to the participant’s T1w image. Linear regression removed mean white-matter and cerebrospinal-fluid signals and 24 motion parameters. We then projected the data to the reconstructed middle cortical surface and smoothed them with a 2-mm FWHM kernel [50].

We used the 100-region Schaefer cortical atlas, which partitions the cortex into visual, sensorimotor, default-mode, dorsal-attention, ventral-attention, limbic, and control networks [51]. Spherical registration mapped the atlas to each participant’s surface [48]. For each scan, we correlated every pair of regional time series to form a functional-connectivity matrix and averaged within-network correlations for each of the seven networks.

### Diffusion MRI processing

We processed diffusion MRI with our infant-specific pipeline [52]. Automated quality control excluded interrupted scans and data with uncorrectable distortion or signal dropout [53].

Usable scans required complete diffusion data in one or two opposing phase-encoding directions, no more than moderate motion or distortion, and at least one QC-passed T1w or T2w image. We corrected signal dropout, pileup, eddy-current distortion, and susceptibility distortion with rotation-invariant contrasts and structural MRI [54], then coregistered diffusion and structural images with ANTs. Visual inspection verified registration, especially in newborns. We fitted diffusion tensors to data with b < 1,500 s/mm^2^ and derived FA, MD, and MO.

We registered FA maps to the ENIGMA FA template in 1-mm MNI152 space and extracted skeletonized values from 34 tracts in the JHU ICBM-DTI-81 atlas [55, 56]. Applying the same deformation fields to MD and MO maps yielded tract-level MD and MO. Quality control included visual checks of eigenvector orientation and FA registration.

### Statistical analyses

#### Permutational multivariate analysis of variance

We quantified beta diversity with Aitchison distances and tested covariate associations with PERMANOVA using adonis2 in the vegan R package [57]. We first tested each birth-related and demographic covariate separately, then entered all covariates into a joint model to estimate their collective association with metagenomic composition.

#### Outlier removal

For each neuroimaging phenotype, we set values at least five median absolute deviations from the median to missing; affected participants could therefore differ across phenotypes. For metagenomic data, we excluded 16 of 991 samples with fewer than 3 million post-QC reads and 20 samples without participant identifiers, leaving 955 samples.

#### Nonlinear developmental trajectories and age adjustment

We removed age trends from neuroimaging phenotypes before association analysis. Generalized additive mixed models in mgcv (R 3.6.0) included a thin-plate spline for age, s(age, bs = ‘tp’, k = 10), and a participant random intercept, s(ID, bs = ‘re’) [58]. Restricted maximum likelihood selected smoothing parameters. Downstream analyses used the age-adjusted residuals.

#### Neuroimaging PC scores

We reduced the dimensions of age-adjusted regional structural and diffusion measures with PCA. After z-scaling residuals, we analyzed (i) cortical volume, surface area, and thickness across 148 Destrieux regions and (ii) tract-average FA, MD, and MO across 34 white-matter tracts (Supplementary Tables S5–S10). An automated elbow criterion selected the retained components from each scree plot ([59]; Supplementary Figures S11 and S12). Downstream microbiome–imaging and CCA models used these PC scores.

#### Microbiome-neuroimaging association analyses

Models adjusted for site, age, age squared, sex, sex-by-age terms, fecal sequencing depth, maternal education, delivery mode, and feeding practice at four months; morphological models also adjusted for total brain volume. The dataset lacked time-varying breastfeeding, recent illness, gastrointestinal symptoms, antibiotic use, and detailed diet, leaving possible residual confounding. We tested microbial features present in at least 10% of participants and controlled FDR within each prespecified microbiome–neuroimaging family.

We used MaAsLin2 to test associations of species, genera, GMMs, GBMs, MetaCyc pathways, and diversity indices with neuroimaging measures [60]. Each generalized linear mixed model included participant as a random effect. The model structure was as follows:

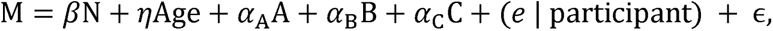

Here, M denotes a microbiome trait, N the neuroimaging measure, A–C the confounding covariates, (e | participant) the participant-specific random effect, and ε the residual error. We tested microbiome features present in at least 10% of samples with non-zero abundance; sample size varied with imaging availability (Supplementary Table S1). Before modeling, we centered-log-ratio transformed species, genera, GMMs, GBMs, and MetaCyc pathways. We z-scaled diversity indices without CLR transformation.

Benjamini–Hochberg FDR correction was applied separately within each prespecified neuroimaging–microbiome family (Supplementary Table S3). Associations with q < 0.05 were considered statistically significant. Associations with 0.05 ≤ q < 0.25 are reported only in the Supplementary Information as exploratory estimates and are not described as significant. Supplementary Excel Table S4 reports summary statistics for the spatially distributed regional analyses.

#### MWAS sensitivity analysis

To test sensitivity to stool–MRI timing, we repeated the MWAS for species, GMMs, GBMs, MetaCyc pathways, and diversity indices after pairing each stool sample with the closest structural, diffusion, or functional MRI observation from the same child within maximum intervals of 15, 20, or 30 days; each stool sample contributed at most one match per imaging modality. Models retained the primary feature filters, transformations, covariates, participant random effect, and family-specific Benjamini–Hochberg correction, with additional adjustment for the signed stool-minus-MRI age difference (in months). Categorical factors retained the default reference-level ordering used in the original MaAsLin2 analysis. For each window, we summarized associations meeting q < 0.05 and compared the effect estimates and 95% confidence intervals for the five primary Figure 5 signals (Supplementary Figure S14 and Supplementary Table S11).

#### Longitudinal slope-covariation analysis

We tested whether participant-specific microbiome and neuroimaging slopes covaried. We retained GMMs, GBMs, species, and MetaCyc features present in at least 10% of participants. Before centered-log-ratio transformation, we replaced zeros with half the smallest positive abundance; we did not CLR-transform diversity measures. Separate growth models for each standardized feature included fixed effects for standardized age, age squared, sex, both age-by-sex terms, site, maternal education, delivery mode, and feeding practice at four months, plus participant-specific random intercepts and age slopes. Microbiome models also adjusted for log sequencing depth. Surface-area and volume models and their PCs also adjusted for total brain volume. The supplied functional-connectivity files lacked a head-motion measure.

We first fitted correlated random intercepts and slopes. If that model was singular, we fitted uncorrelated random effects; if slope variance remained singular, we estimated within-participant slopes from covariate-adjusted residuals by ordinary least squares. Participants required at least two distinct ages, and each feature model and slope correlation required at least 10 eligible participants. We retained a paired slope only when both values satisfied |slope − median slope| < 5 × MAD. Pearson correlations and Fisher-transformed 95% confidence intervals quantified slope covariation. We controlled FDR within each prespecified microbiome–neuroimaging family and used q < 0.05 as the significance threshold. Post hoc age-stratified analyses divided participants into mean-age tertiles and controlled FDR across six tests. These correlations assess coordinated change, not temporal precedence. We fitted growth models with lme4.

#### Longitudinal CCA-score covariation and cross-lag analysis

We applied frozen CCA1 and CCA2 weights to eligible longitudinal observations and modeled each standardized score separately by domain. Fixed effects included standardized age, age squared, sex, both age-by-sex terms, site, maternal education, delivery mode, and feeding practice at four months. Microbiome models also included standardized log sequencing depth; neuroimaging models included standardized total brain volume. We fitted correlated participant-specific random intercepts and age slopes by restricted maximum likelihood: score = fixed effects + (1 + age | participant). If the fit failed or was singular at 10□□, we fitted uncorrelated random effects: score = fixed effects + (1 | participant) + (0 + age | participant).

When slope variance remained singular or negligible, we regressed the standardized score on the fixed covariates and fitted adjusted score against standardized age separately for each participant by ordinary least squares. All four domain-by-axis models used this fallback.

Participants required at least two distinct ages and a positive age span; each score model required at least 20 observations and 10 repeatedly measured participants. We matched microbiome and neuroimaging slopes by participant and retained pairs only when both slopes satisfied |slope − median slope| < 5 × MAD. Pearson correlations, Fisher-transformed 95% confidence intervals, and equivalent slope-on-slope regressions quantified association. Tests required at least 10 matched participants. FDR was controlled across the six CCA slope-correlation tests.

We defined signed lag as neuroimaging age minus microbiome age. Positive windows of 1–3, 3–6, and 6–12 months tested earlier microbiome against later neuroimaging scores; negative windows of −3 to −1, −6 to −3, and −12 to −6 months tested the reverse direction. We formed all within-participant candidate pairs, then matched visits one-to-one within each window, prioritizing lags closest to the midpoint. No visit was reused within a model. Complete pairs also had to satisfy |score − median score| < 5 × MAD for both exposure and outcome. Each model required at least 25 pairs from 20 participants.

For each axis, window, and direction, we regressed the standardized later score on the standardized earlier score. Covariates included standardized response age, age squared, sex, both response-age-by-sex terms, site, maternal education, delivery mode, feeding practice at four months, standardized log sequencing depth, standardized total brain volume, and standardized absolute lag. The primary autoregressive model also included the latest outcome-domain score observed at or before the exposure age. In neuroimaging-to-microbiome models, the coefficient estimates whether earlier neuroimaging CCA predicts later microbiome CCA beyond the latest prior microbiome score and measured covariates.

We first fitted each cross-lag model by maximum likelihood with a participant random intercept. If its variance was singular at 10□□, we refitted the same fixed effects by ordinary least squares and used participant-clustered HC1 standard errors. These fallback t tests and 95% confidence intervals used participants minus one denominator degrees of freedom. We applied Benjamini–Hochberg correction separately by analysis type and axis across the six directional windows and defined significance as q < 0.05. Mixed models used lme4 and lmerTest; clustered inference used sandwich and lmtest.

#### Canonical correlation analysis

To balance dimensionality and interpretation, CCA included GMMs, GBMs, species, and diversity indices (Supplementary Table S4). We selected species for taxonomic resolution, GMMs and GBMs for interpretable functional summaries, and diversity indices for community structure. After outlier removal, we retained 33 GMMs, 10 GBMs, and 30 species that together captured more than 80% of mean abundance within each class. These 73 abundance features plus five diversity indices formed the 78-variable microbiome block. The 22-variable brain block contained three PCs each for cortical area, thickness, and volume; two PCs each for FA, MD, and MO; and connectivity within seven functional networks (Supplementary Table S2).

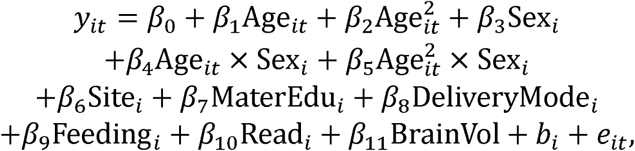

Here, β terms denote fixed effects, b□ the participant random intercept, and e□□ residual error. We extracted participant-level random-intercept estimates, which capture trait variation not explained by measured covariates, and z-scaled them for multivariate analysis. Restricting the data to complete multimodal observations yielded matrices of 107 participants × 22 brain traits and 107 participants × 78 microbiome traits.

We fitted ridge-regularized CCA to the adjusted microbiome (X) and brain (Y) matrices with mixOmics 3.22 [61] in R 4.4.0. Ten-fold cross-validation selected regularization parameters λ = 0.20, λ = 0.95. We compared observed correlations with 1,000 label permutations and assessed generalization in 1,000 repeated 75%/25% train/test splits. Each split estimated loadings only in the training set before applying them to held-out participants. We interpreted a mode only when its correlation exceeded the 95th percentile of the permutation distribution and remained positive in more than 85% of held-out splits. These safeguards reduce but do not eliminate overfitting; independent replication remains necessary. The regularized objective is shown below.

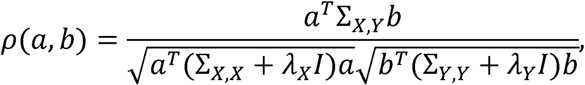

Here, ΣXX and ΣYY denote the within-block covariance matrices, ΣXY denotes the cross-block covariance matrix. The solution yields ordered orthogonal canonical variates; subsequent modes capture progressively smaller components of shared covariance.

#### Sparse partial least-squares analysis

We fitted sparse canonical PLS with mixOmics [61] to the same standardized residual matrices used for rCCA (107 participants × 78 microbiome features and 107 participants × 22 neuroimaging features). The canonical-mode model estimated 20 paired latent components.

Although the theoretical maximum number of PLS components equals the number of neuroimaging features (n = 22), extracting components at the full rank of the neuroimaging matrix risks numerical instability in the terminal components; we therefore set a conservative ceiling of 20 components. We correlated every native PLS score with the fixed CCA1 and CCA2 scores separately in the microbiome and neuroimaging blocks. Joint CCA–PLS correspondence was the mean of these two block-specific correlations. PLS signs were aligned only after model fitting, using both blocks simultaneously, because latent-axis orientation is arbitrary. Feature contributions were Pearson correlations between each input variable and its block-specific latent score; shared direction was evaluated for features with |r| ≥ 0.20 in both the CCA and sign-aligned PLS mode.

We assessed out-of-sample behavior in 1,000 participant-level random 90%/10% training–test splits using the same random seed as the CCA evaluation. Centering, scaling, the sparse PLS fit, and sequential deflation coefficients were estimated using only the training set and then applied to the test set. Because component order can change after refitting, reference PLS1, PLS2, and PLS5 were matched jointly and one-to-one to the refitted components by maximizing the mean absolute cosine similarity of their microbiome and neuroimaging loading vectors. We calculated the Pearson correlation between the matched microbiome and neuroimaging scores separately in the training and test sets and summarized their means and the percentage of held-out correlations greater than zero. We also evaluated associations of microbiome PLS1 and PLS2 scores with auditory–visual volume difference, and of microbiome PLS5 scores with ventral-attention network connectivity, using Pearson correlations, two-sided P values, and 95% confidence intervals.

## Data availability

MWAS and CCA summary statistics are available in Supplementary Excel Tables S1–S4. Individual-level BCP demographic and MRI data are available through the NIMH Data Archive (https://nda.nih.gov) under NDA accession DOI 10.15154/nshn-2b72. Individual-level microbiota data from the BCP-Enriched study are available through the European Nucleotide Archive (https://www.ebi.ac.uk/ena/browser/home) under accession PRJEB73364 (secondary accession ERP158163).

## Code availability

Analysis code written in R and Python is available at Zenodo with synthetic example data (https://doi.org/10.5281/zenodo.16267242), and MRI processing used iBEAT V2.0 (http://www.ibeat.cloud/). The repository includes a README, an input-file manifest (including reads.txt where applicable), environment or package-version information, and nonidentifiable example inputs sufficient to demonstrate each workflow step; controlled individual-level data remain available only through the repositories listed above.

## Acknowledgements

We thank Dr. Norbert Sprenger for helpful conversations. We thank the individuals represented in the UNC/UMN Baby Connectome Project Consortium for their participation and the research teams for their work in collecting, processing, and disseminating these datasets for analysis. This work was supported by NIH Grants U01MH110274 (W.L., J.T.E., J.P.), 3U01MH110274-03S1 (W.L., J.T.E., J.P.), 1U01DA055344 (W.L.), R01MH104324 (J.T.E.), R01MH104324-S1 (J.T.E.), R01MH125479 (P.T.Y.), R01EB008374 (P.T.Y.), R01MH133836 (P.T.Y.), RF1AG082938 (H.Z.), R01AG085581 (H.Z.), R01AR082684 (H.Z.), R01MH136055 (T.L., H.Z.), R21HD120911 (T.L.), and K01AG095286 (T.L.). The funders had no role in study design, data collection and analysis, decision to publish or preparation of the manuscript.

## Author contributions

T.L., Y.Y., and W.L. designed research, Y.Y. processed the microbiome data and performed PERMANOVA, trajectory fitting and MWAS, T.L. performed CCA, PLS, covariation, cross-lag, sensitivity analyses and all visualizations, Z.W., W.Y., T.L., K.M.H., G.L., P.T.Y., L.W. processed the imaging data, W.L., S.H.C., B.R.H. performed research investigations, T.L.,Y.Y., S.H.C., B.R.H., W.Y., N.S., J.T.E., S.A., H.Z., and W.L. wrote the paper. T.L. and Y.Y. contributed equally to the work.

## Additional information

### Supplementary Information

Supplementary Information is available for this paper.

