## Supplementary_Materials for "Microbes on the Mind: Multi-modal Neuroimaging Reveals Gut-Brain Axes in Infants"

Weili Lin, Ph.D.

Biomedical Research Imaging Center CB#7513

University of North Carolina at Chapel Hill

This supplementary file includes:

Supplementary Figures S1–S14

Supplementary Tables S1–S11

Supplementary Text includes the infant study protocol, supplementary methods, analyses, results, and interpretations.

Other Supplementary Material for this manuscript includes the following:

(separate Excel files; Supplementary Data)

Supplementary Excel Tables S1–S4 (.xlsx)

Supplementary Figures

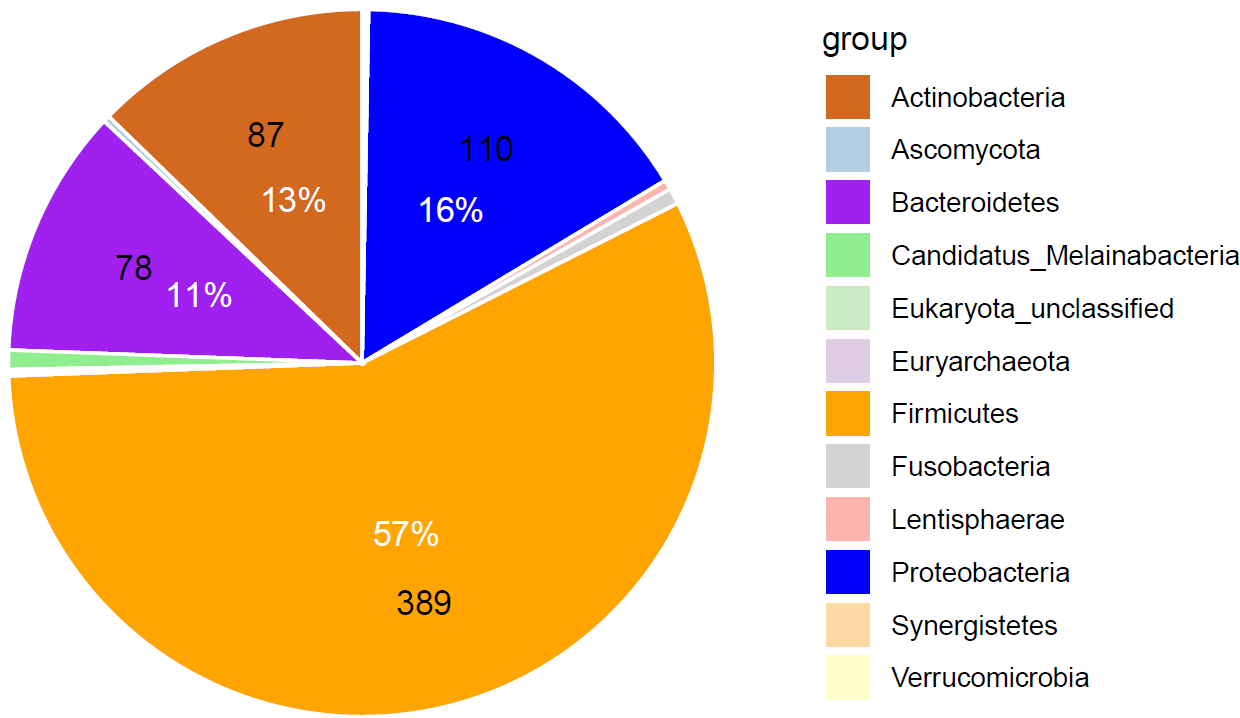

**Supplementary Figure S1. Numbers (black) and proportions (white) of species assigned to each phylum.**

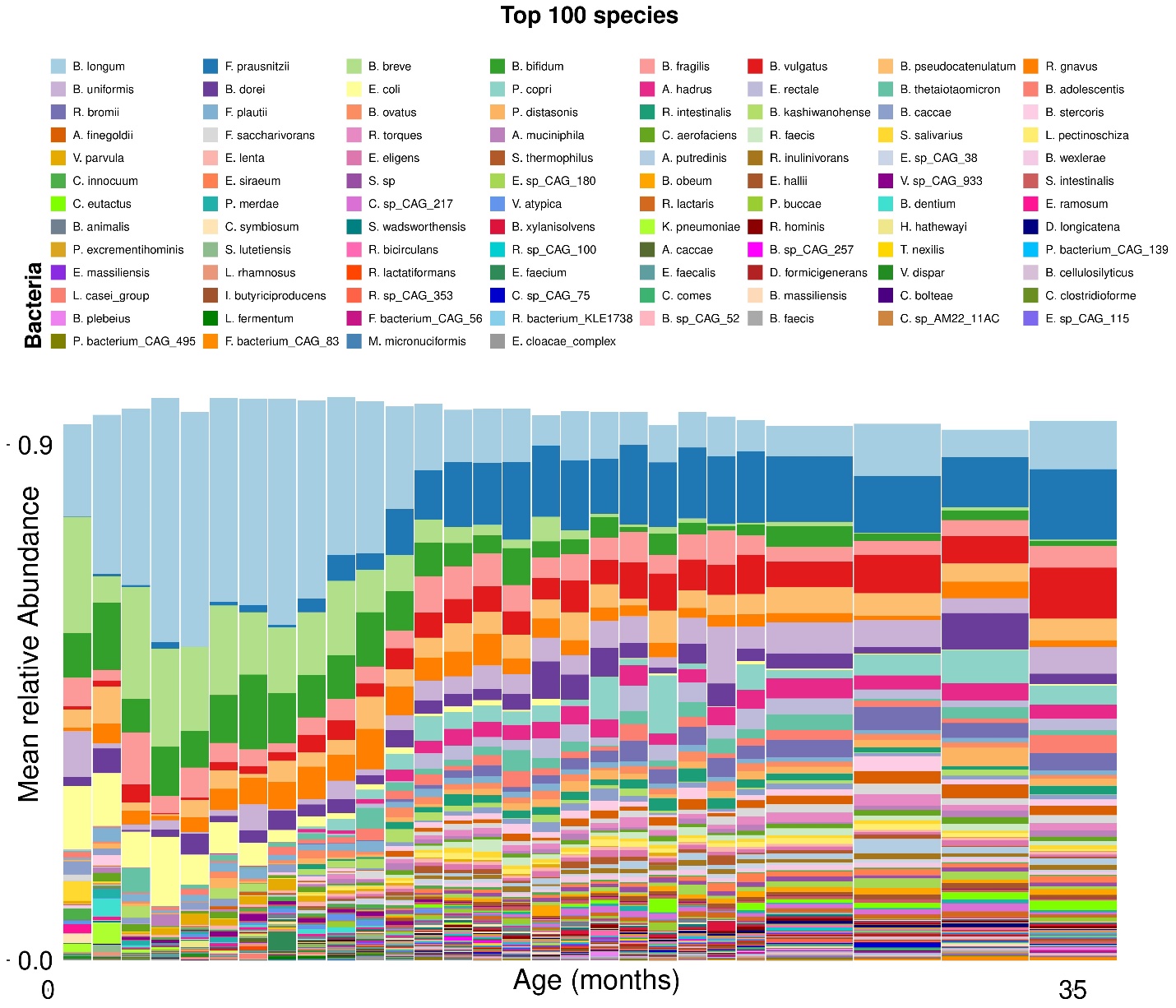

**Supplementary Figure S2. Temporal distribution of the 100 most abundant species during the first three years of life. This expanded display complements the top-10 panel in main Figure 3. Abundances are normalized within each age bin, and the y-axis shows mean relative abundance. Monthly summaries are shown from 0–24 months and at three-month intervals thereafter.**

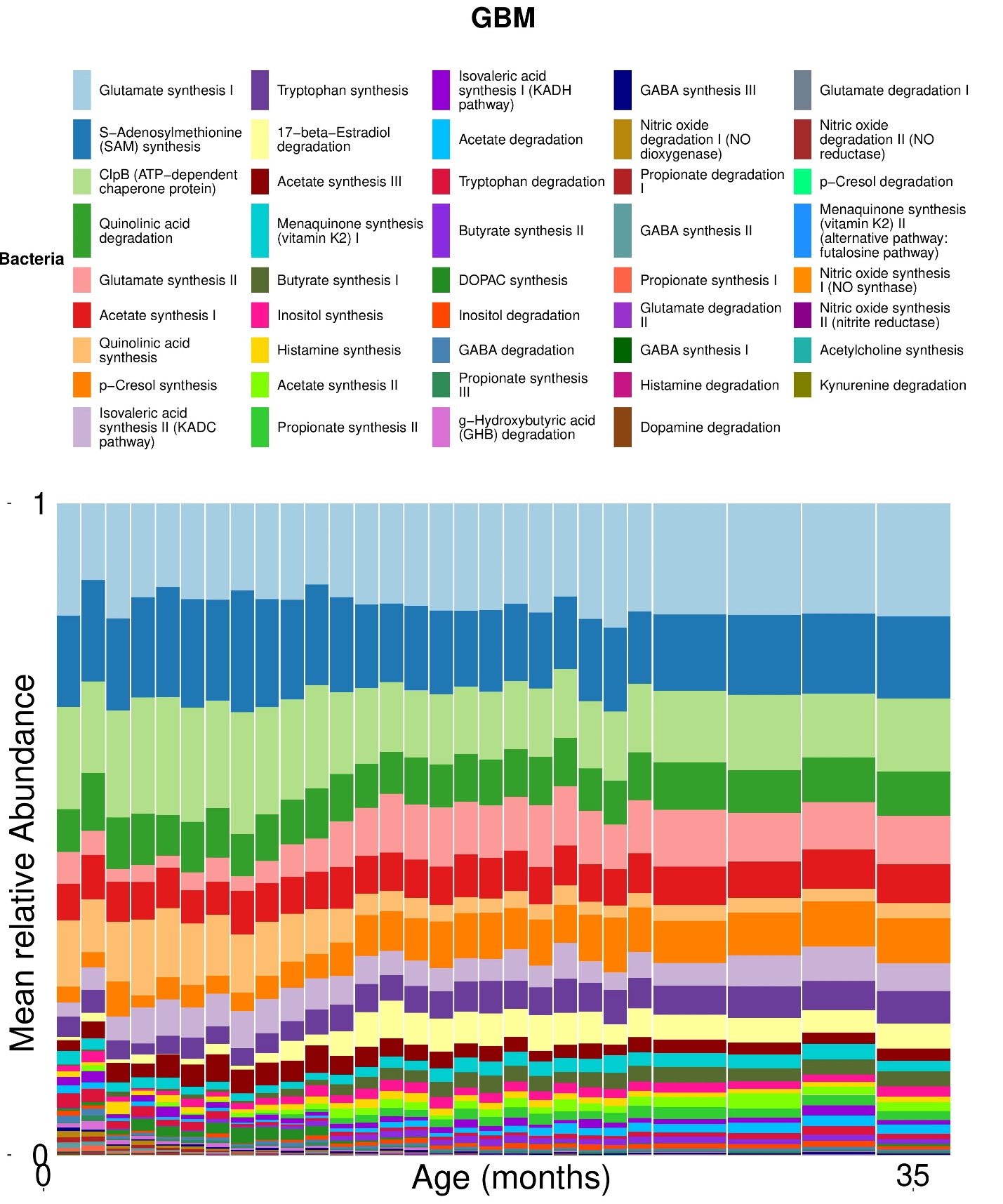

**Supplementary Figure S3. Temporal distribution of the relative abundance of all 44 GBMs during the first three years of life. Monthly summaries are shown from 0–24 months and at three-month intervals thereafter.**

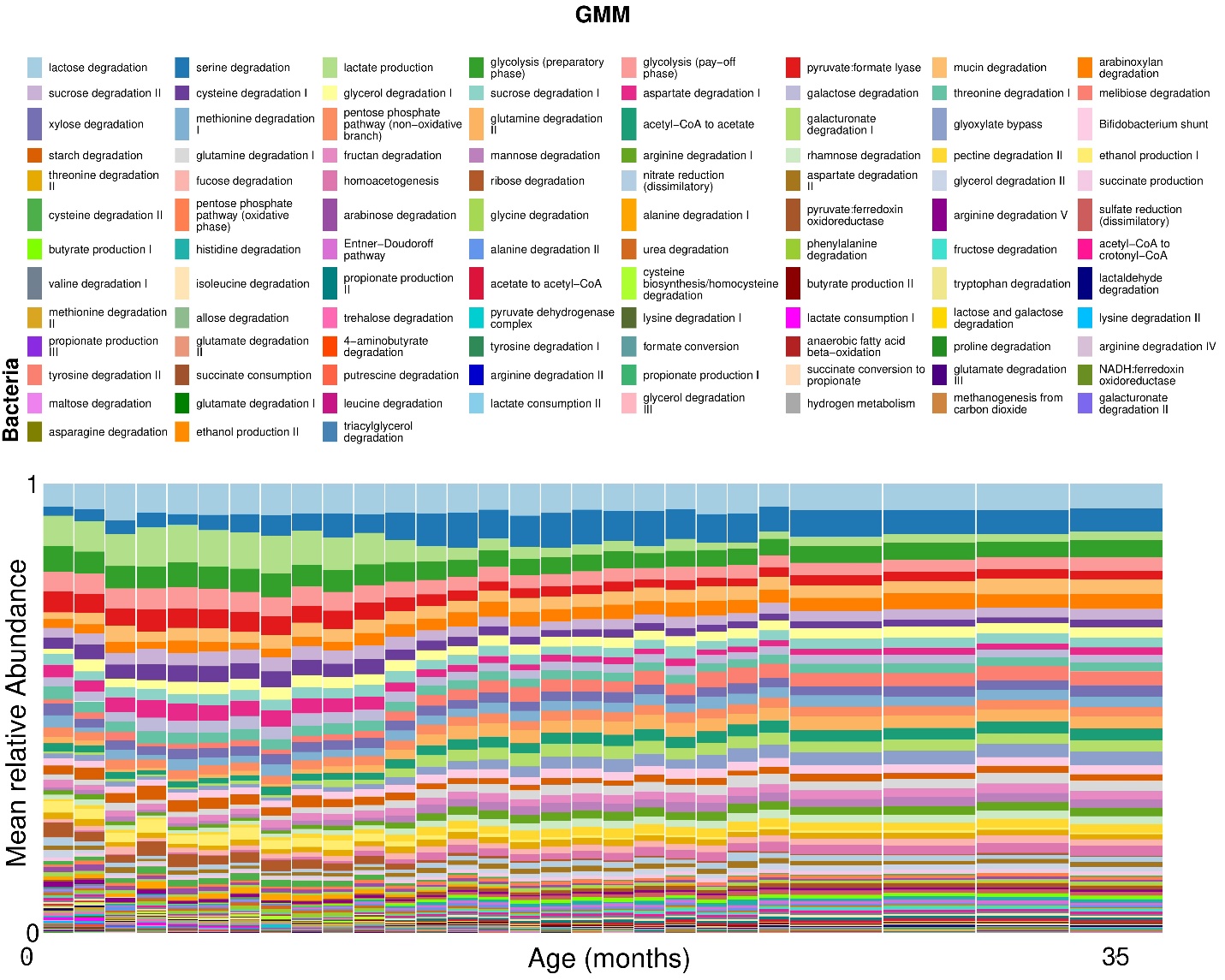

**Supplementary Figure S4. Temporal distribution of the relative abundance of all 99 GMMs during the first three years of life. Monthly summaries are shown from 0–24 months and at three-month intervals thereafter.**

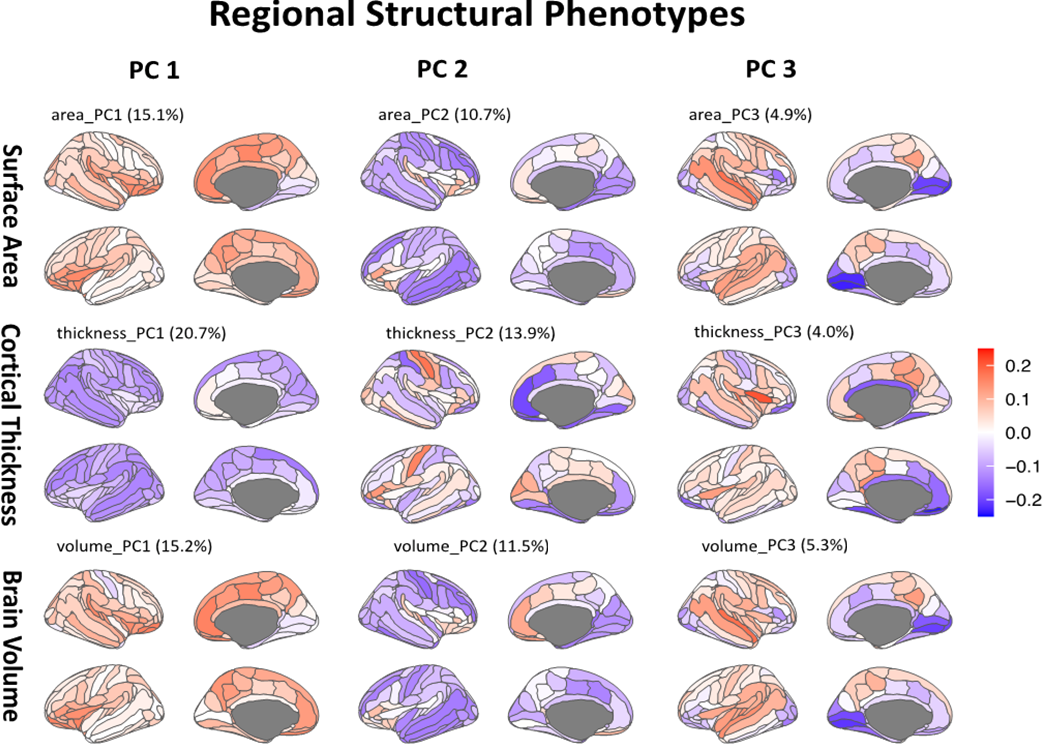

**Supplementary Figure S5. Loading coefficients for the first three principal components (PCs) of regional cortical structural measures across 148 regions. Rows correspond to surface area, cortical thickness, and cortical volume, and columns correspond to PC1–PC3; percentages in parentheses indicate variance explained. Red denotes positive loadings and blue denotes negative loadings.**

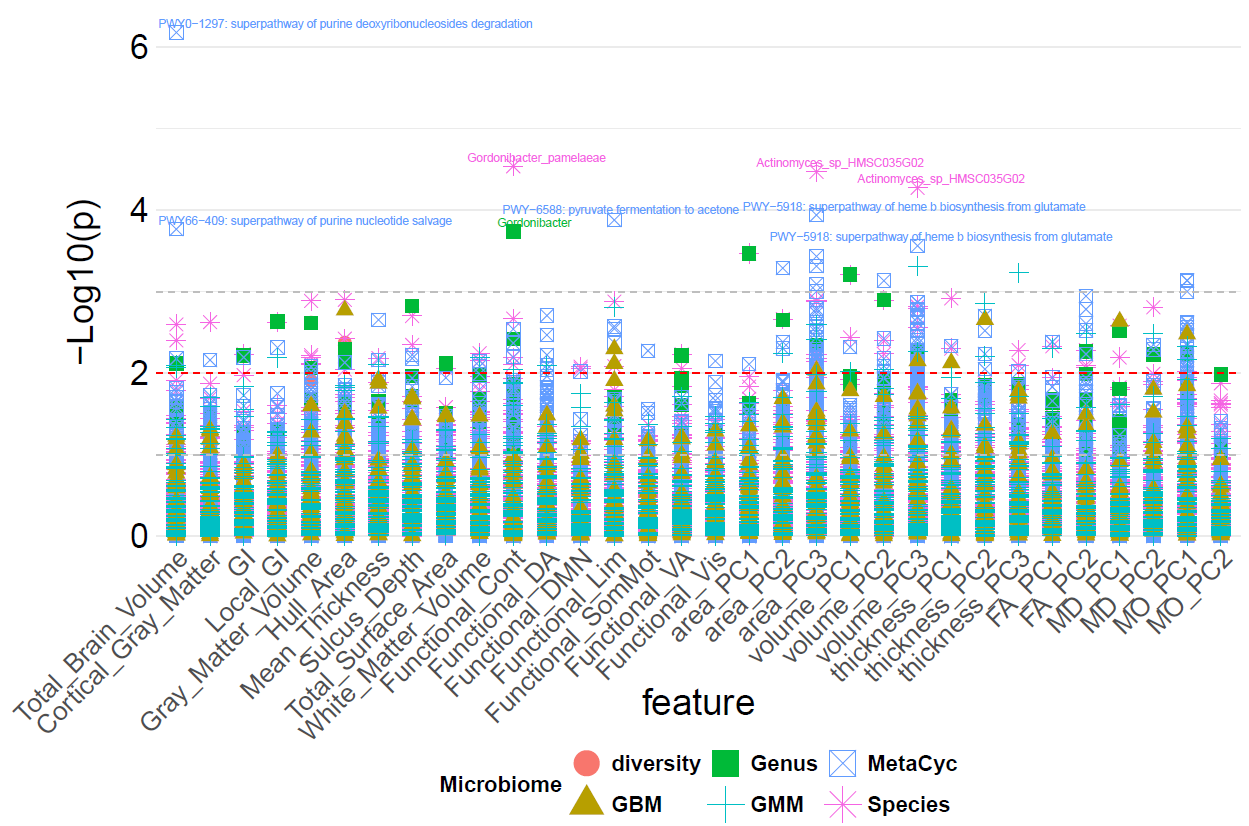

**Supplementary Figure S6. Manhattan plot illustrating associations between neuroimaging phenotypes and gut microbiome traits.** The y-axis shows the –log₁₀ of raw p-values for each microbiome–neuroimaging association. Each column on the x-axis represents a distinct neuroimaging phenotype, and each point corresponds to a microbiome trait, distinguished by shape.

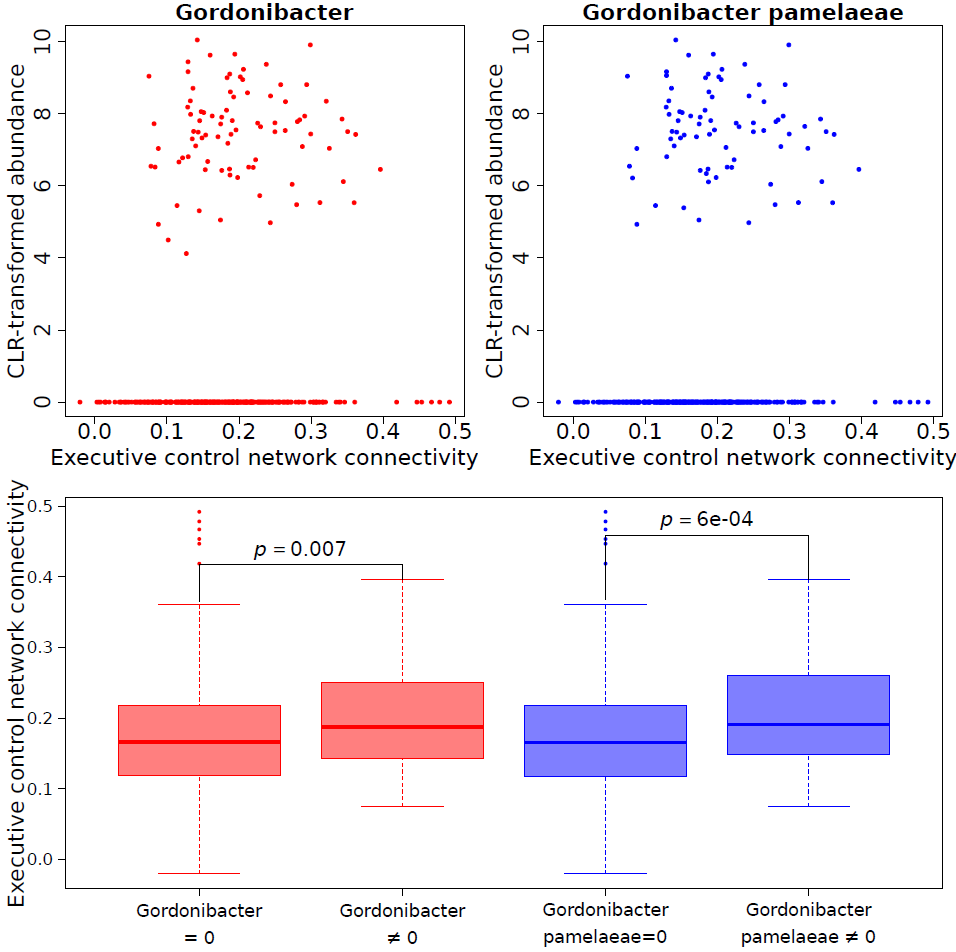

**Supplementary Figure S7. Scatterplots (upper panels) and boxplots (lower panels) of CLR-transformed *Gordonibacter pamelaeae* and *Gordonibacter* abundance in relation to executive-control network connectivity. Only the species-level association met q < 0.05; the genus-level association is exploratory.**

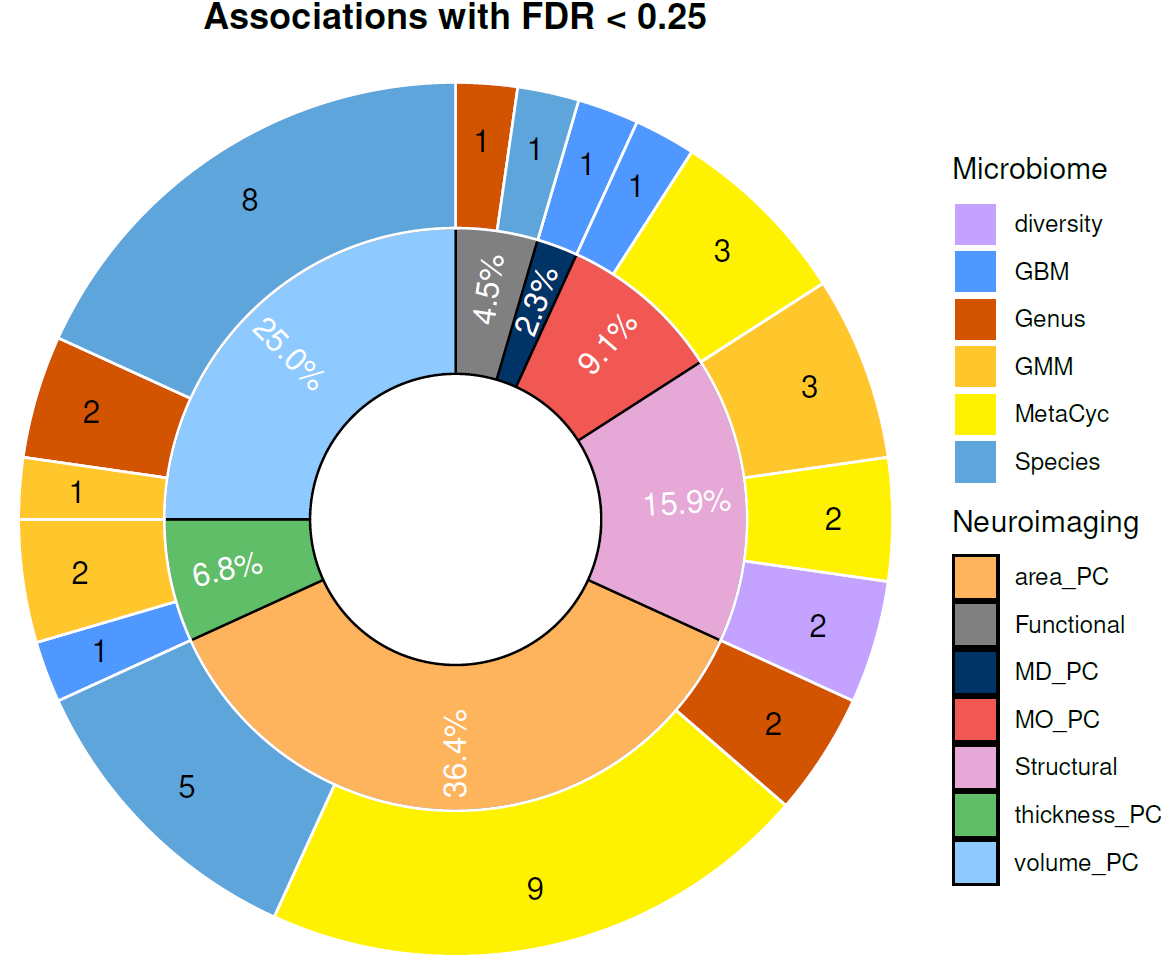

**Supplementary Figure S8. Distribution of associations with q < 0.25. Associations with 0.05 ≤ q < 0.25 are retained for hypothesis generation and are not treated as statistically significant. The inner circle groups associations by neuroimaging phenotype, and the outer annotations show counts by microbial feature class.**

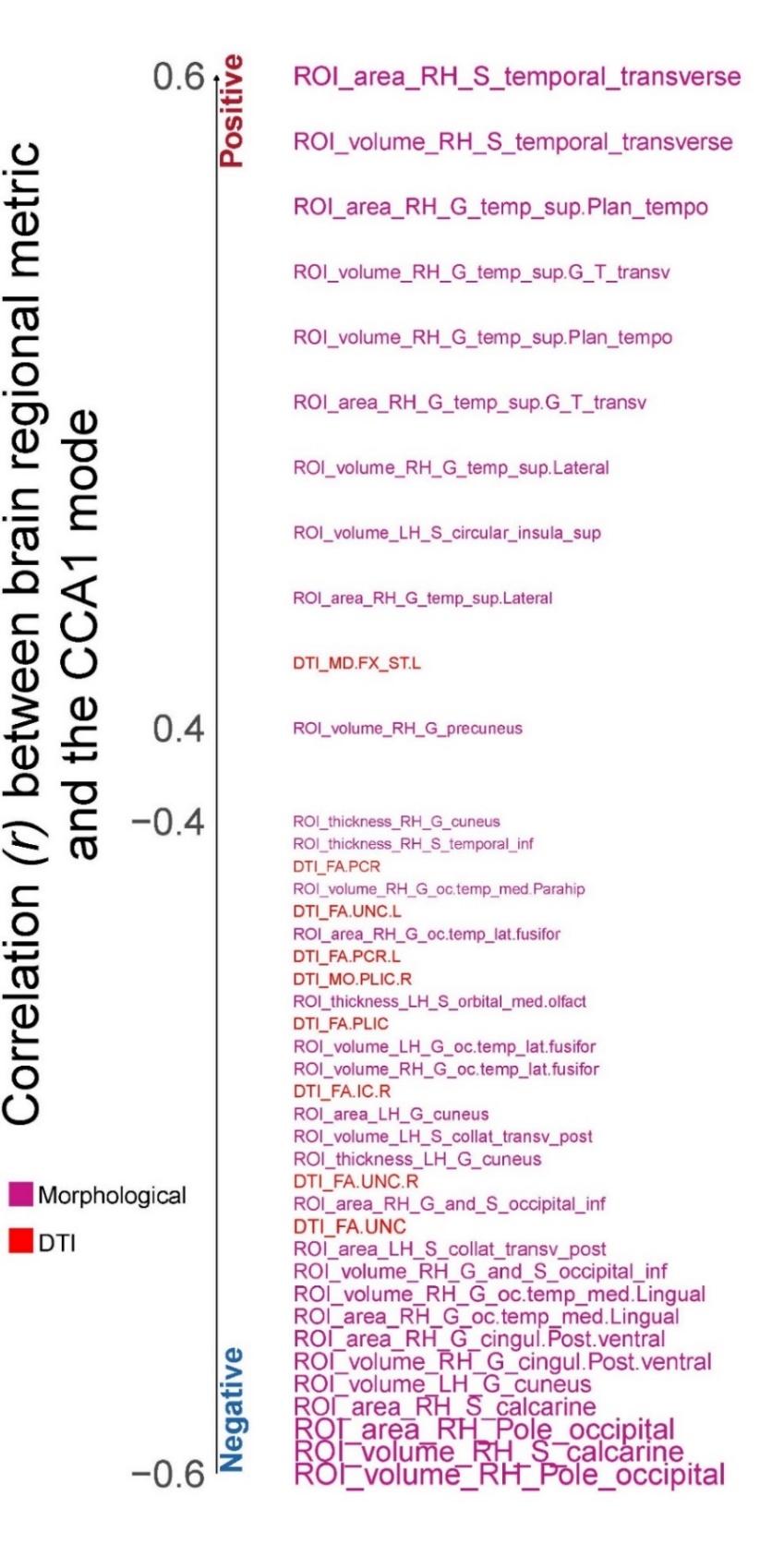

**Supplementary Figure S9. Pearson correlations between the first canonical variate and regional brain measures. Regional measures include cortical volume, surface area, and thickness across 148 regions of interest and diffusion parameters (fractional anisotropy, mean diffusivity, and mode of diffusivity) across 34 white-matter tracts.**

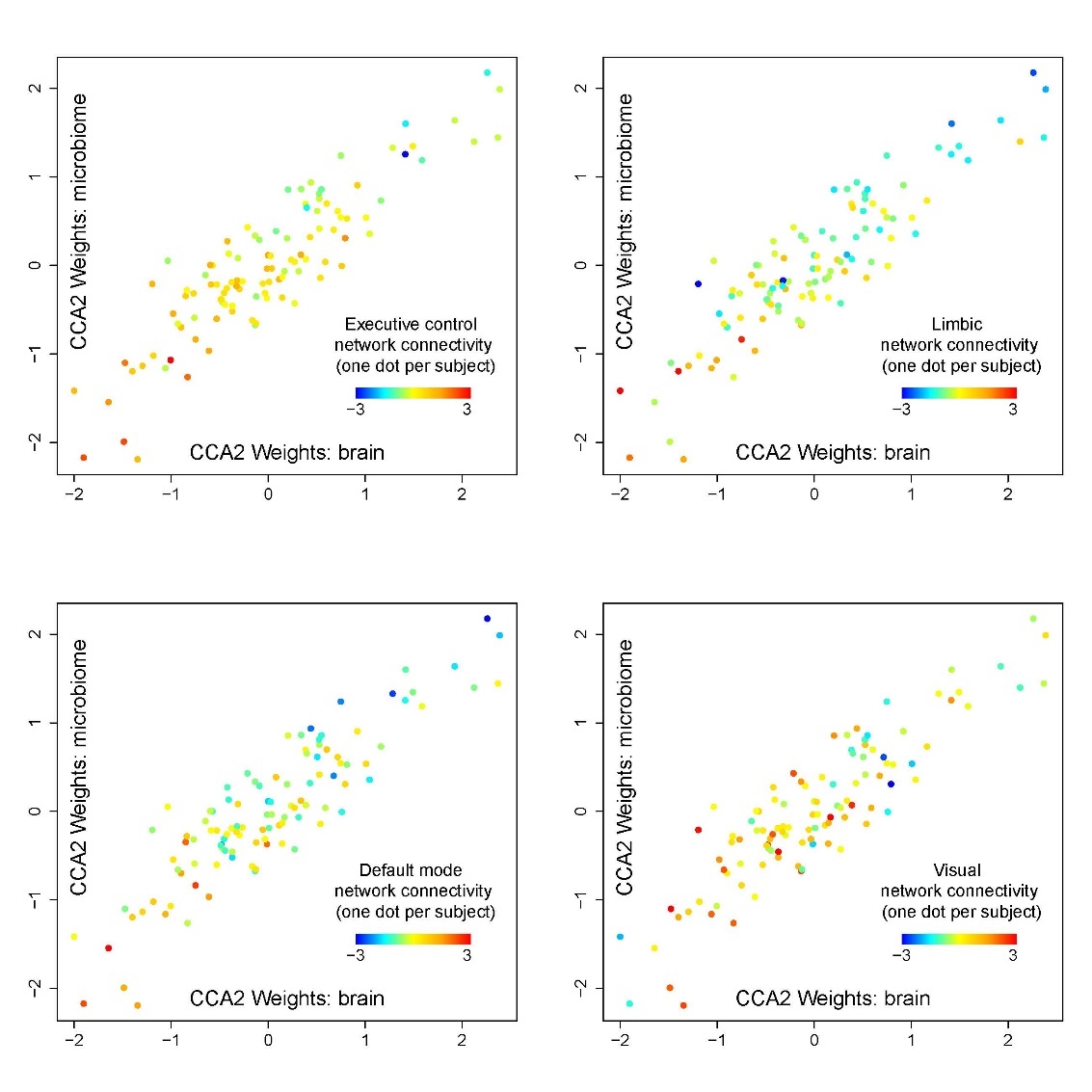

**Supplementary Figure S10. Scatterplots relating CCA2 scores to within-network connectivity in executive-control, limbic, default-mode, and visual networks. Each point represents one participant. The panels display the observations underlying the multivariate functional pattern and should not be interpreted as evidence that microbial variation causes differences in connectivity.**

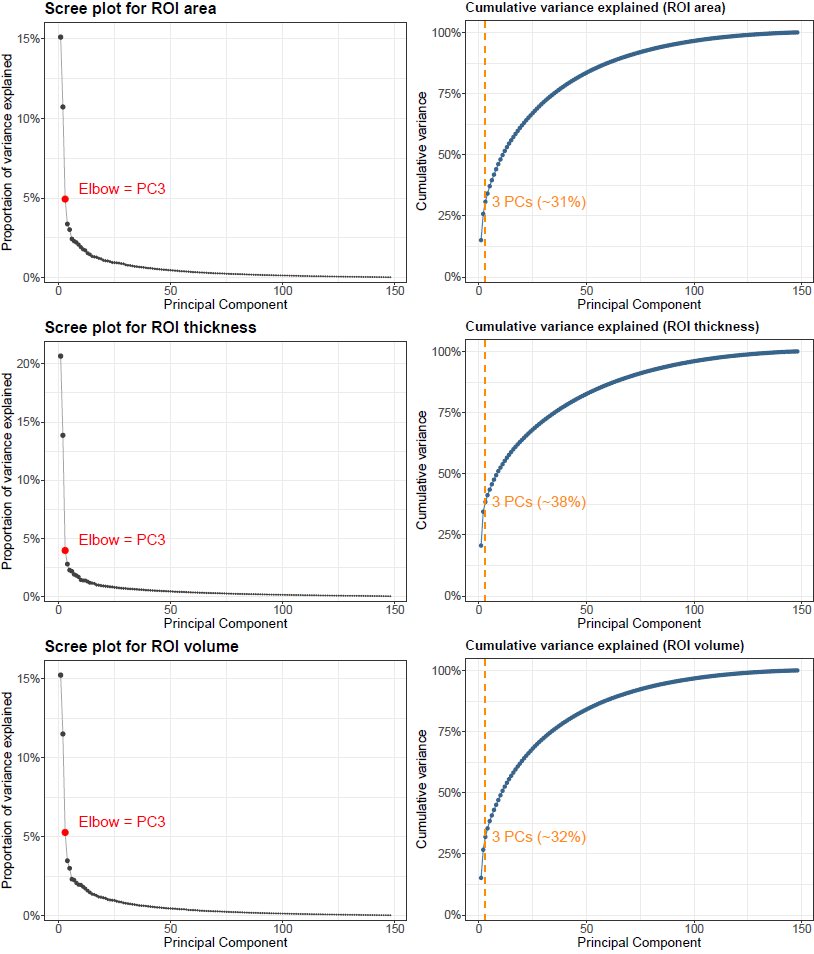

**Supplementary Figure S11. Selection of principal components (PCs) for regional cortical structural phenotypes. Left, scree plots showing the proportion of variance explained by each PC for surface area, cortical thickness, and volume across 148 cortical regions. Right, cumulative variance explained for the corresponding measures. Red dots and blue lines indicate the PCs retained for subsequent analyses.**

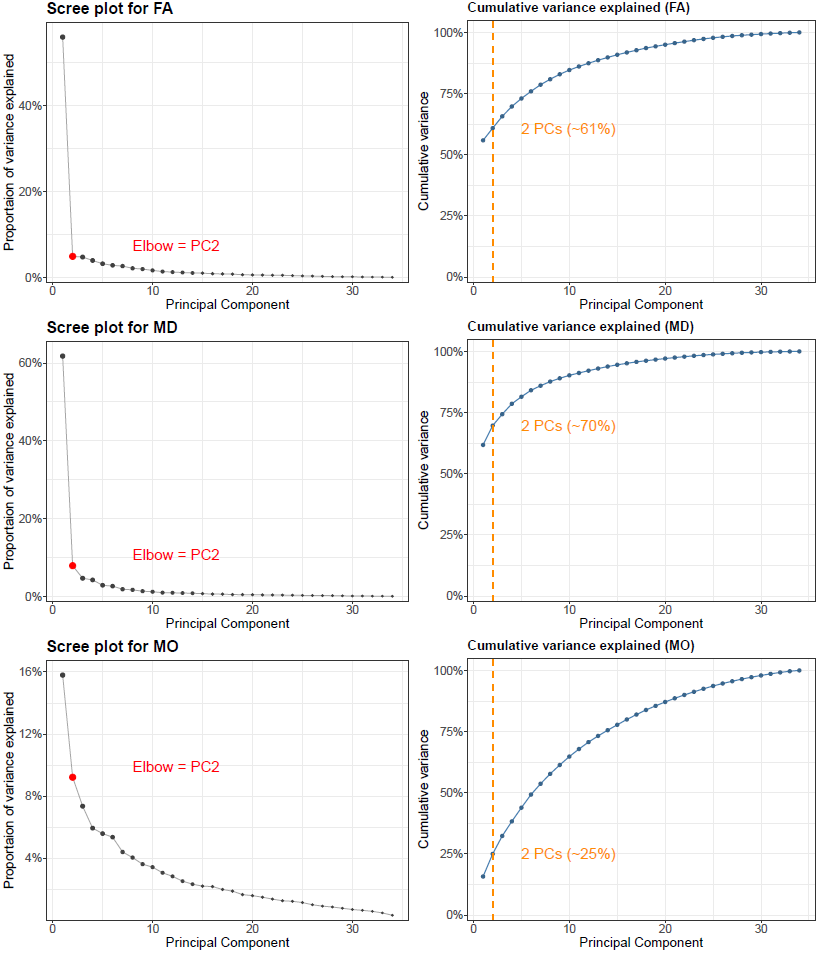

**Supplementary Figure S12. Selection of principal components (PCs) for regional white-matter diffusion measures across 34 tracts. Left, scree plots for fractional anisotropy (FA), mean diffusivity (MD), and mode of diffusivity (MO). Right, cumulative variance explained for the corresponding measures. Red dots and blue lines indicate the PCs retained for subsequent analyses.**

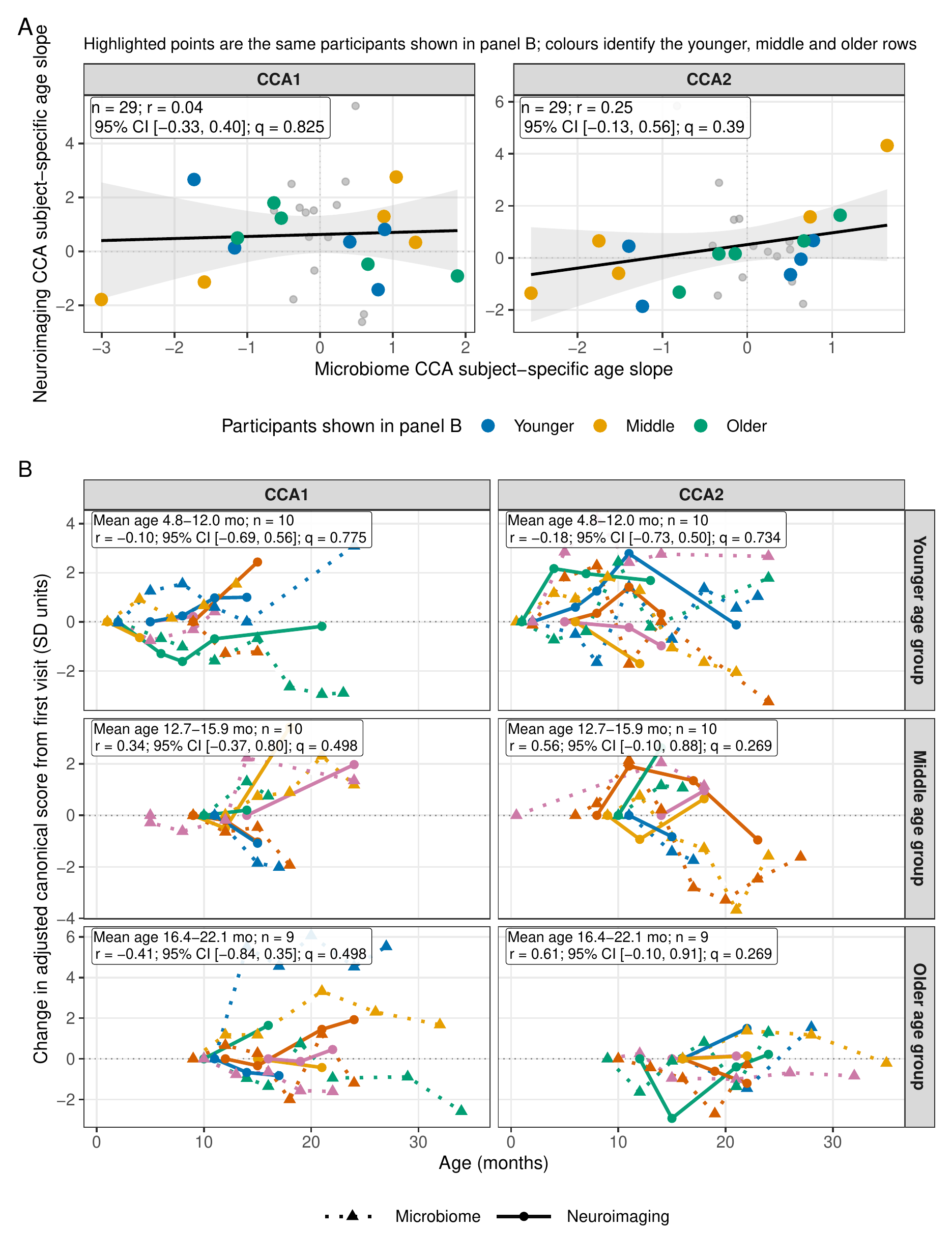

**Supplementary Figure S13. Longitudinal covariation of microbiome and neuroimaging CCA scores.** A, Pearson correlations between covariate-adjusted participant-specific age slopes for matched microbiome and neuroimaging canonical scores. All 29 participants passed the five-MAD filter; neither CCA1 nor CCA2 was significant after FDR correction. B, Adjusted longitudinal score changes for five representative participants in each tertile of mean longitudinal age. Dotted lines denote microbiome scores and solid lines denote neuroimaging scores. Text boxes summarize correlations for all participants in each age group, not only the displayed examples; q values are FDR-adjusted across the six age-stratified tests. Frozen CCA weights were applied to longitudinal observations before score modeling. MAD, median absolute deviation; FDR, false discovery rate.

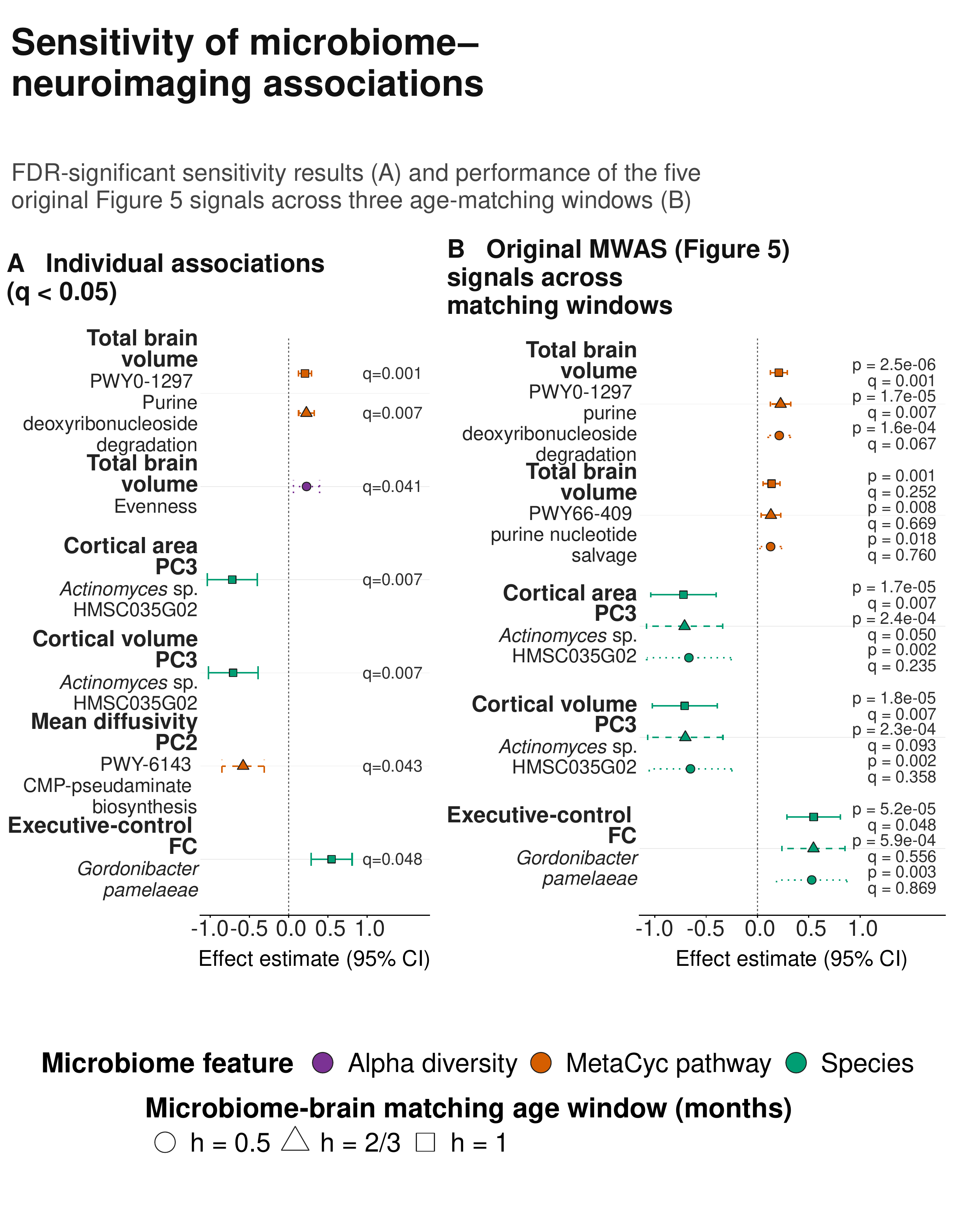

**Supplementary Figure S14. Sensitivity analysis of microbiome-wide associations across stool–MRI matching windows.** A, MWAS associations meeting q < 0.05 in the sensitivity analyses. B, Effect estimates and 95% confidence intervals for the five associations highlighted in main Figure 5 across matching windows of 15 days (h = 0.5 month), 20 days (h = 2/3 month), and 30 days (h = 1 month). Point shape and confidence-interval line pattern identify the matching window; labels report nominal p values and FDR-adjusted q values. All sensitivity models included the signed stool-minus-MRI age difference in months as an additional covariate. Narrower windows reduced analytic sample sizes, as summarized in Supplementary Table S11.

Supplementary Tables

**Supplementary Table S1. Number of microbiome traits tested for each neuroimaging phenotype. Microbiome–neuroimaging association analyses were restricted to microbial traits present in at least 10% of participants; consequently, the number of tested traits differed across neuroimaging phenotype sets.**

| **Neuroimaging/Microbiome** | **Diversity** | **Species** | **Genus** | **MetaCyc** | **GBM** | **GMM** |
| --- | --- | --- | --- | --- | --- | --- |
| Structural Phenotypes/Structural PCs | 5 | 135 | 61 | 371 | 31 | 77 |
| Diffusivity PCs | 5 | 141 | 61 | 363 | 31 | 75 |
| Functional Phenotypes | 5 | 134 | 61 | 371 | 31 | 77 |

**Supplementary Table S2. Neuroimaging features included in each analysis category.**

| **Neuroimaging** | **Features** |
| --- | --- |
| Global (whole brain) structural phenotypes | Total Brain Volume, Cortical Gray Matter, Gyrification Index, Gray Matter Volume, Hull Area, Local Gyrification Index, Mean Thickness, Sulcus Depth, Total Surface Area, and White Matter Volume |
| Brain Regional Structural phenotypes and numbers of PCs | {Surface Area, Volume, Thickness} * {PC1, PC2, PC3} |
| Functional Networks | Executive Control Network, Dorsal Attention Network, Default Mode Network, Limbic Network, Somatosensory-Motor Network, Salience Network, and Visual Network |
| Diffusion parameters and number of PCs | {Fractional Anisotropy, Mean Diffusivity, Mode of Anisotropy} * {PC1, PC2} |

PC, principal component.

**Supplementary Table S3. Number of tests included in FDR correction for each neuroimaging phenotype and microbiome feature class. The numbers of microbial traits available after matching to neuroimaging phenotypes are reported in Supplementary Table S1. FA, fractional anisotropy; MD, mean diffusivity; MO, mode of diffusivity; PC, principal component.**

| **Neuroimaging/Microbiome** | **Diversity** | **Species** | **Genus** | **MetaCyc** | **GBM** | **GMM** |
| --- | --- | --- | --- | --- | --- | --- |
| All Volume | 5 | 135 | 61 | 371 | 31 | 77 |
| Other Overall Structural | 5*9 | 135*9 | 61*9 | 371*9 | 31*9 | 77*9 |
| Cortical Area PCs | 5*3 | 135*3 | 61*3 | 371*3 | 31*3 | 77*3 |
| Cortical Thickness PCs | 5*3 | 135*3 | 61*3 | 371*3 | 31*3 | 77*3 |
| Cortical Volume PCs | 5*3 | 135*3 | 61*3 | 371*3 | 31*3 | 77*3 |
| FA PCs | 5*2 | 141*2 | 61*2 | 363*2 | 31*2 | 75*2 |
| MD PCs | 5*2 | 141*2 | 61*2 | 363*2 | 31*2 | 75*2 |
| MO PCs | 5*2 | 141*2 | 61*2 | 363*2 | 31*2 | 75*2 |
| Functional Phenotypes | 5*7 | 134*7 | 61*7 | 371*7 | 31*7 | 77*7 |

**Supplementary Table S4. Microbiome traits included in the canonical correlation analysis. PC, principal component.**

| **Microbiome** | **Members** |
| --- | --- |
| Diversity (5) | Shannon, evenness, richness, beta diversity (PC1 and PC2) |
| GBM (10) | Glutamate.synthesis.I,S.Adenosylmethionine..SAM..synthesis,ClpB..ATP.dependent.chaperone.protein.,Quinolinic.acid.degradation,Glutamate.synthesis.II,Acetate.synthesis.I,Quinolinic.acid.synthesis,p.Cresol.synthesis,Isovaleric.acid.synthesis.II..KADC.pathway.,Tryptophan.synthesis |
| GMM (33) | Lactose degradation, serine degradation, lactate production, glycolysis..preparatory.phase, glycolysis..pay.off.phase., pyruvate formate lyase, mucin degradation, arabinoxylan degradation, sucrose degradation II, cysteine degradation I, glycerol degradation I, sucrose degradation I, aspartate degradation I, galactose degradation, threonine degradation I, melibiose.degradation,xylose.degradation,methionine.degradation.I,pentose.phosphate.pathway..non.oxidative.branch.,glutamine.degradation.II,acetyl.CoA.to.acetate,galacturonate.degradation.I,glyoxylate.bypass,Bifidobacterium.shunt,starch.degradation,glutamine.degradation.I,fructan.degradation,mannose.degradation,arginine.degradation.I,rhamnose.degradation,pectine.degradation.II,ethanol.production.I,threonine.degradation.II |
| Species (30) | *Bifidobacterium longum*, *Faecalibacterium prausnitzii*, *Bifidobacterium breve*, *Bifidobacterium bifidum*, *Bacteroides fragilis*, *Bacteroides vulgatus*, *Bifidobacterium pseudocatenulatum*, *Ruminococcus gnavus*, *Bacteroides uniformis*, *Bacteroides dorei*, *Escherichia coli*, *Prevotella copri*, *Anaerostipes hadrus*, *Eubacterium rectale*, *Bacteroides thetaiotaomicron*, *Bifidobacterium adolescentis*, *Ruminococcus bromii*, *Flavonifractor plautii*, *Bacteroides ovatus*, *Parabacteroides distasonis*, *Roseburia intestinalis*, *Bifidobacterium kashiwanohense*, *Bacteroides caccae*, *Bacteroides stercoris*, *Alistipes finegoldii*, *Fusicatenibacter saccharivorans*, *Ruminococcus torques*, *Akkermansia muciniphila*, *Collinsella aerofaciens*, *Roseburia faecis* |

**Supplementary Table S5. The 30 cortical regions with the largest absolute loading coefficients for cortical volume in each of the first three principal components (PCs).**

|  | **PC1** | | **PC2** | | **PC3** | |
| --- | --- | --- | --- | --- | --- | --- |
| **structure** | **LH_PC1** | **RH_PC1** | **LH_PC2** | **RH_PC2** | **LH_PC3** | **RH_PC3** |
| G_Ins_lg_and_S_cent_ins | 0.0119425 |  |  |  |  | 0.01042391 |
| G_and_S_cingul.Ant | 0.01251017 | 0.01540748 |  |  |  |  |
| G_and_S_cingul.Mid.Ant |  | 0.01271755 | -0.0128154 |  |  |  |
| G_and_S_cingul.Mid.Post |  | 0.01491999 | -0.0116323 |  |  |  |
| G_and_S_frontomargin |  |  | -0.0109588 |  |  |  |
| G_and_S_occipital_inf |  |  |  |  | -0.0143743 | -0.0135146 |
| G_and_S_paracentral |  |  |  |  |  |  |
| G_and_S_subcentral |  |  |  |  |  | 0.01063263 |
| G_and_S_transv_frontopol |  |  |  |  |  |  |
| G_cingul.Post.dorsal |  |  |  |  |  |  |
| G_cingul.Post.ventral |  |  |  | -0.0149707 |  |  |
| G_cuneus |  |  |  |  | -0.0135669 |  |
| G_front_inf.Opercular |  |  |  |  |  |  |
| G_front_inf.Orbital |  |  |  |  |  |  |
| G_front_inf.Triangul | 0.01237519 |  |  |  |  | -0.0113741 |
| G_front_middle |  |  | -0.0132652 | -0.0147409 |  |  |
| G_front_sup |  | 0.0116727 |  |  |  |  |
| G_insular_short | 0.01545317 | 0.01433367 |  |  |  |  |
| G_oc.temp_lat.fusifor |  |  |  | -0.0125724 | -0.0121866 |  |
| G_oc.temp_med.Lingual |  |  |  |  | -0.0214908 | -0.0205268 |
| G_oc.temp_med.Parahip |  |  |  | -0.0135204 |  |  |
| G_occipital_middle |  |  | -0.0129532 |  |  |  |
| G_occipital_sup |  |  |  |  | -0.0114892 |  |
| G_orbital | 0.01538443 | 0.01571932 |  |  |  |  |
| G_pariet_inf.Angular |  |  |  |  |  |  |
| G_pariet_inf.Supramar |  |  |  |  | 0.01034843 |  |
| G_parietal_sup |  |  |  |  |  |  |
| G_postcentral |  |  |  | -0.0129972 |  |  |
| G_precentral |  |  |  | -0.0126335 |  |  |
| G_precuneus | 0.01325422 | 0.01248216 |  |  |  |  |
| G_rectus | 0.0145098 | 0.01426982 |  | 0.01187635 |  |  |
| G_subcallosal |  |  |  |  |  |  |
| G_temp_sup.G_T_transv |  | 0.01287986 |  |  | 0.01126493 |  |
| G_temp_sup.Lateral |  |  | -0.0123299 |  | 0.01430212 | 0.01789492 |
| G_temp_sup.Plan_polar | 0.01135817 | 0.01255241 |  |  |  |  |
| G_temp_sup.Plan_tempo |  | 0.01494986 |  |  | 0.01043035 |  |
| G_temporal_inf |  |  | -0.0129008 | -0.0147133 |  |  |
| G_temporal_middle |  |  | -0.0137757 |  |  |  |
| Lat_Fis.ant.Horizont |  | 0.01199682 |  |  |  |  |
| Lat_Fis.ant.Vertical |  |  |  |  |  |  |
| Lat_Fis.post |  |  |  |  |  |  |
| Pole_occipital |  |  |  |  | -0.0155992 | -0.0196313 |
| Pole_temporal |  |  | -0.0138358 | -0.0144292 |  |  |
| S_calcarine |  |  |  | -0.0118077 | -0.0195231 | -0.0179168 |
| S_central |  |  | -0.0127501 | -0.0156159 |  |  |
| S_cingul.Marginalis |  | 0.01220527 |  |  |  |  |
| S_circular_insula_ant | 0.01199778 | 0.01235811 |  |  |  |  |
| S_circular_insula_inf |  |  | -0.0125452 |  | 0.01264636 |  |
| S_circular_insula_sup | 0.01389301 |  |  |  | 0.01112899 |  |
| S_collat_transv_ant |  |  |  | -0.0125276 |  |  |
| S_collat_transv_post |  |  |  |  | -0.017161 | -0.0157396 |
| S_front_inf |  |  |  | -0.0125122 |  |  |
| S_front_middle |  |  |  |  |  |  |
| S_front_sup |  |  |  |  |  |  |
| S_interm_prim.Jensen |  |  |  |  |  |  |
| S_intrapariet_and_P_trans |  |  |  |  |  |  |
| S_oc.temp_lat |  |  |  | -0.0121159 |  |  |
| S_oc.temp_med_and_Lingual |  |  |  | -0.0109772 | -0.0160983 |  |
| S_oc_middle_and_Lunatus |  |  |  |  |  | -0.0106895 |
| S_oc_sup_and_transversal |  |  |  |  |  |  |
| S_occipital_ant |  |  | -0.0110774 |  |  |  |
| S_orbital.H_Shaped | 0.01111692 | 0.01571632 |  |  |  |  |
| S_orbital_lateral |  |  |  |  |  |  |
| S_orbital_med.olfact | 0.01364006 | 0.01355619 |  |  |  |  |
| S_parieto_occipital |  |  |  | -0.0109557 |  |  |
| S_pericallosal |  |  |  |  |  |  |
| S_postcentral |  |  |  |  |  |  |
| S_precentral.inf.part |  |  |  |  |  |  |
| S_precentral.sup.part |  |  |  |  |  |  |
| S_suborbital |  |  |  |  | -0.0102683 |  |
| S_subparietal | 0.01247274 |  |  |  |  | 0.01256516 |
| S_temporal_inf |  |  | -0.0109303 |  |  |  |
| S_temporal_sup |  |  | -0.0137235 |  |  | 0.01227809 |
| S_temporal_transverse |  | 0.01187959 |  |  | 0.01387685 | 0.01138016 |

**Supplementary Table S6. The 30 cortical regions with the largest absolute loading coefficients for cortical surface area in each of the first three principal components (PCs).**

|  | **PC1** | | **PC2** | | **PC3** | |
| --- | --- | --- | --- | --- | --- | --- |
| **structure** | **LH_PC1** | **RH_PC1** | **LH_PC2** | **RH_PC2** | **LH_PC3** | **RH_PC3** |
| G_Ins_lg_and_S_cent_ins |  |  |  |  |  |  |
| G_and_S_cingul.Ant |  | 0.014699798 |  |  |  |  |
| G_and_S_cingul.Mid.Ant |  |  | -0.012364109 |  |  |  |
| G_and_S_cingul.Mid.Post |  | 0.014909176 | -0.011420524 |  |  |  |
| G_and_S_frontomargin |  |  |  |  |  |  |
| G_and_S_occipital_inf |  |  |  |  | -0.011361229 | -0.015066388 |
| G_and_S_paracentral |  |  |  |  |  |  |
| G_and_S_subcentral |  |  |  |  |  | 0.010373829 |
| G_and_S_transv_frontopol |  |  |  |  |  |  |
| G_cingul.Post.dorsal |  |  |  |  |  |  |
| G_cingul.Post.ventral |  |  |  | -0.015550595 |  |  |
| G_cuneus |  |  |  |  | -0.016364698 |  |
| G_front_inf.Opercular |  |  |  |  |  |  |
| G_front_inf.Orbital |  |  |  |  |  |  |
| G_front_inf.Triangul | 0.01256785 |  | 0.011164901 |  |  | -0.015251263 |
| G_front_middle |  |  | -0.014197655 | -0.015235278 |  |  |
| G_front_sup | 0.011349484 | 0.012676553 |  |  |  |  |
| G_insular_short | 0.014456317 | 0.014723005 |  |  |  |  |
| G_oc.temp_lat.fusifor |  |  |  | -0.013413802 |  |  |
| G_oc.temp_med.Lingual |  |  |  | -0.011487128 | -0.023990883 | -0.019799239 |
| G_oc.temp_med.Parahip |  |  |  | -0.014488383 |  |  |
| G_occipital_middle |  |  | -0.013385292 |  |  |  |
| G_occipital_sup |  |  |  |  | -0.012594036 |  |
| G_orbital | 0.014772952 | 0.014168362 |  |  |  |  |
| G_pariet_inf.Angular |  |  |  |  |  |  |
| G_pariet_inf.Supramar |  |  |  |  | 0.010771949 |  |
| G_parietal_sup |  |  |  |  |  |  |
| G_postcentral |  |  |  |  |  |  |
| G_precentral |  |  |  | -0.011562505 |  |  |
| G_precuneus | 0.012863392 | 0.011522148 |  |  |  |  |
| G_rectus | 0.013874851 |  |  |  |  |  |
| G_subcallosal |  |  |  |  |  |  |
| G_temp_sup.G_T_transv |  | 0.012924831 |  |  | 0.011094928 |  |
| G_temp_sup.Lateral |  |  | -0.012372911 |  | 0.012661357 | 0.018210053 |
| G_temp_sup.Plan_polar | 0.012192351 | 0.012070227 |  |  |  |  |
| G_temp_sup.Plan_tempo |  | 0.01481194 |  |  |  |  |
| G_temporal_inf |  |  | -0.014164814 | -0.015746847 |  |  |
| G_temporal_middle |  |  | -0.014973035 |  |  |  |
| Lat_Fis.ant.Horizont | 0.012741276 | 0.012808523 |  |  |  |  |
| Lat_Fis.ant.Vertical | 0.011995664 |  |  |  |  |  |
| Lat_Fis.post |  |  |  |  | 0.010801125 |  |
| Pole_occipital |  |  |  |  | -0.019406487 | -0.023008397 |
| Pole_temporal |  |  | -0.013780992 | -0.014143369 |  |  |
| S_calcarine |  |  |  | -0.011833307 | -0.023169356 | -0.020923996 |
| S_central |  |  |  | -0.012493949 |  |  |
| S_cingul.Marginalis |  | 0.012462271 |  |  |  |  |
| S_circular_insula_ant | 0.012401322 | 0.014348547 |  |  | -0.010940102 |  |
| S_circular_insula_inf |  |  | -0.012574539 | -0.011201105 | 0.012618014 | 0.010810964 |
| S_circular_insula_sup | 0.014677752 | 0.011795584 |  |  |  |  |
| S_collat_transv_ant |  |  |  | -0.012304957 |  |  |
| S_collat_transv_post |  |  |  |  | -0.016855914 | -0.016305716 |
| S_front_inf |  |  |  | -0.013588613 |  |  |
| S_front_middle |  |  |  |  |  |  |
| S_front_sup |  |  | -0.011757141 |  |  |  |
| S_interm_prim.Jensen |  |  |  |  |  |  |
| S_intrapariet_and_P_trans |  |  |  |  |  |  |
| S_oc.temp_lat |  |  | -0.01128004 | -0.011957167 |  |  |
| S_oc.temp_med_and_Lingual |  |  |  | -0.012075816 | -0.010986882 |  |
| S_oc_middle_and_Lunatus |  |  |  |  | -0.010709659 | -0.010590562 |
| S_oc_sup_and_transversal |  |  |  |  |  |  |
| S_occipital_ant |  |  | -0.012225205 |  |  |  |
| S_orbital.H_Shaped | 0.011466918 | 0.015457819 |  |  |  |  |
| S_orbital_lateral |  |  |  |  |  |  |
| S_orbital_med.olfact | 0.01414192 | 0.014292757 |  |  |  |  |
| S_parieto_occipital |  |  |  |  |  |  |
| S_pericallosal |  |  |  |  |  |  |
| S_postcentral |  |  |  | -0.012488891 |  |  |
| S_precentral.inf.part |  |  |  |  |  |  |
| S_precentral.sup.part |  |  |  |  |  |  |
| S_suborbital |  |  |  |  |  |  |
| S_subparietal | 0.012419603 |  |  |  |  | 0.012459936 |
| S_temporal_inf |  |  |  |  |  |  |
| S_temporal_sup |  |  | -0.013968651 |  | 0.010568298 | 0.014091827 |
| S_temporal_transverse |  | 0.012789123 |  |  | 0.010236857 | 0.012276827 |

**Supplementary Table S7. The 30 cortical regions with the largest absolute loading coefficients for cortical thickness in each of the first three principal components (PCs).**

|  | **PC1** | | **PC2** | | **PC3** | |
| --- | --- | --- | --- | --- | --- | --- |
| **structure** | **LH_PC1** | **RH_PC1** | **LH_PC2** | **RH_PC2** | **LH_PC3** | **RH_PC3** |
| G_Ins_lg_and_S_cent_ins |  |  |  |  |  | 0.013235461 |
| G_and_S_cingul.Ant |  |  | -0.011160417 | -0.0196678 | -0.017042652 |  |
| G_and_S_cingul.Mid.Ant |  |  |  | -0.017394544 | -0.012834696 |  |
| G_and_S_cingul.Mid.Post |  |  |  |  |  |  |
| G_and_S_frontomargin |  |  |  |  |  |  |
| G_and_S_occipital_inf |  |  | -0.014250281 |  |  |  |
| G_and_S_paracentral |  |  |  |  |  |  |
| G_and_S_subcentral |  | -0.010198465 |  |  |  |  |
| G_and_S_transv_frontopol |  |  |  |  |  |  |
| G_cingul.Post.dorsal |  |  |  |  | 0.014293877 | 0.014049495 |
| G_cingul.Post.ventral |  |  | -0.014216216 |  | 0.013888698 | -0.013965312 |
| G_cuneus |  | -0.01004169 | 0.012605137 |  |  |  |
| G_front_inf.Opercular |  |  |  |  |  |  |
| G_front_inf.Orbital |  |  |  |  |  |  |
| G_front_inf.Triangul |  | -0.009971244 | 0.013341339 |  |  |  |
| G_front_middle | -0.012162116 | -0.011504008 |  |  |  |  |
| G_front_sup | -0.012818218 | -0.010283025 |  |  |  |  |
| G_insular_short |  |  |  |  |  |  |
| G_oc.temp_lat.fusifor |  |  | -0.014215017 |  | -0.019002382 |  |
| G_oc.temp_med.Lingual |  |  | -0.013473071 | -0.016479859 |  |  |
| G_oc.temp_med.Parahip |  |  | -0.014062001 | -0.01755894 | -0.012329564 |  |
| G_occipital_middle | -0.010182701 | -0.010375725 |  |  |  |  |
| G_occipital_sup | -0.009563735 | -0.009509245 |  | 0.011155716 |  |  |
| G_orbital |  |  | -0.012561483 | -0.013070972 | -0.018452878 | -0.017365709 |
| G_pariet_inf.Angular | -0.011080696 | -0.012162854 |  |  |  |  |
| G_pariet_inf.Supramar | -0.011867165 |  |  |  |  |  |
| G_parietal_sup | -0.009963116 |  |  | -0.012581211 |  |  |
| G_postcentral |  |  |  | 0.010529168 |  |  |
| G_precentral | -0.010552667 |  |  | 0.010526711 |  |  |
| G_precuneus | -0.009544139 | -0.009615315 |  |  |  |  |
| G_rectus |  |  | -0.016511899 | -0.019273584 | -0.020899324 |  |
| G_subcallosal |  |  |  |  | -0.014073773 | 0.01725479 |
| G_temp_sup.G_T_transv |  |  |  |  |  | 0.017283797 |
| G_temp_sup.Lateral |  | -0.009665386 |  |  |  |  |
| G_temp_sup.Plan_polar |  |  |  |  |  |  |
| G_temp_sup.Plan_tempo |  |  |  |  |  |  |
| G_temporal_inf |  |  |  |  |  |  |
| G_temporal_middle | -0.010855651 | -0.010951638 |  |  |  |  |
| Lat_Fis.ant.Horizont |  |  | 0.012237116 |  |  |  |
| Lat_Fis.ant.Vertical |  |  |  |  |  |  |
| Lat_Fis.post |  |  |  |  |  | 0.011657193 |
| Pole_occipital |  |  |  |  |  |  |
| Pole_temporal |  |  |  |  |  |  |
| S_calcarine |  |  |  |  |  |  |
| S_central |  |  | 0.015726133 | 0.017421779 |  |  |
| S_cingul.Marginalis |  |  |  |  |  | 0.01276389 |
| S_circular_insula_ant |  |  |  |  |  |  |
| S_circular_insula_inf |  |  |  |  |  |  |
| S_circular_insula_sup |  |  |  |  | 0.013347066 | 0.022081954 |
| S_collat_transv_ant |  |  |  |  | -0.016151651 |  |
| S_collat_transv_post |  |  | -0.014402492 |  |  |  |
| S_front_inf | -0.011585963 | -0.009916406 |  |  |  |  |
| S_front_middle |  |  |  |  |  |  |
| S_front_sup |  |  |  |  |  |  |
| S_interm_prim.Jensen |  | -0.010169641 |  |  |  |  |
| S_intrapariet_and_P_trans | -0.01052884 |  |  |  |  |  |
| S_oc.temp_lat |  |  |  |  |  |  |
| S_oc.temp_med_and_Lingual |  |  | -0.014321044 | -0.012517407 | -0.021108893 | -0.018350379 |
| S_oc_middle_and_Lunatus |  |  |  |  |  |  |
| S_oc_sup_and_transversal |  |  |  |  |  |  |
| S_occipital_ant |  |  |  |  |  |  |
| S_orbital.H_Shaped |  |  |  |  | -0.014121393 | -0.013726696 |
| S_orbital_lateral |  |  | 0.01297027 |  |  |  |
| S_orbital_med.olfact |  |  |  |  | -0.0224471 |  |
| S_parieto_occipital |  | -0.009838639 |  |  |  |  |
| S_pericallosal |  |  |  |  | -0.016017149 | -0.018630591 |
| S_postcentral |  |  |  | -0.016327591 |  |  |
| S_precentral.inf.part | -0.009791964 |  |  | -0.010525656 |  |  |
| S_precentral.sup.part |  |  |  |  |  |  |
| S_suborbital |  |  |  | -0.016265519 | -0.025920002 |  |
| S_subparietal |  |  |  |  |  | 0.013590734 |
| S_temporal_inf | -0.009477638 |  |  |  |  | -0.012953496 |
| S_temporal_sup | -0.011859389 | -0.010801689 |  |  |  |  |
| S_temporal_transverse |  |  |  |  |  |  |

**Supplementary Table S8. The tract-based loading coefficients of fractional anisotropy (FA) for the first two principal components (PCs).**

| **ID** | **PC1** | **PC2** |
| --- | --- | --- |
| ACR.L | 0.03123514 | 0.06443862 |
| ACR.R | 0.03129736 | 0.06957414 |
| ALIC.L | 0.03267249 | 0.02280103 |
| ALIC.R | 0.03193969 | 0.00834682 |
| CGC.L | 0.03157357 | 0.02582866 |
| CGC.R | 0.03003441 | 0.00963379 |
| CGH.L | 0.02653183 | -0.0551253 |
| CGH.R | 0.02695432 | -0.0573194 |
| CST.L | 0.0247938 | -0.0049128 |
| CST.R | 0.02738875 | -0.023255 |
| EC.L | 0.03440464 | 0.00574962 |
| EC.R | 0.03344877 | 0.00082509 |
| FX_ST.L | 0.02974699 | -0.0298135 |
| FX_ST.R | 0.02556021 | -0.0437966 |
| IFO.L | 0.02184715 | 0.01857774 |
| IFO.R | 0.02299096 | 0.01137693 |
| PCR.L | 0.03048054 | -0.0082676 |
| PCR.R | 0.03366563 | -0.0175625 |
| PLIC.L | 0.02749343 | 0.00575305 |
| PLIC.R | 0.02400909 | 0.0193899 |
| PTR.L | 0.03238549 | -0.0139945 |
| PTR.R | 0.03041175 | -0.0403178 |
| RLIC.L | 0.03238716 | 0.01223853 |
| RLIC.R | 0.02922552 | 0.01251959 |
| SCR.L | 0.03330728 | 0.03727986 |
| SCR.R | 0.03407619 | 0.03750756 |
| SFO.L | 0.03059511 | 0.04019591 |
| SFO.R | 0.02981715 | 0.01419124 |
| SLF.L | 0.03361989 | 0.00812034 |
| SLF.R | 0.03283538 | 0.01251325 |
| SS.L | 0.03139131 | 0.01120175 |
| SS.R | 0.03085947 | -0.0226807 |
| UNC.L | 0.02175727 | -0.1098411 |
| UNC.R | 0.01926226 | -0.1250498 |

**Supplementary Table S9. The tract-based loading coefficients of mean diffusivity (MD) for the first two principal components (PCs).**

| **ID** | **PC1** | **PC2** |
| --- | --- | --- |
| ACR.L | 0.03379625 | 0.00698134 |
| ACR.R | 0.0335927 | 0.00739026 |
| ALIC.L | 0.03171435 | -0.039409 |
| ALIC.R | 0.03156568 | -0.0369053 |
| CGC.L | 0.03325178 | -0.0108582 |
| CGC.R | 0.03360206 | -0.0112863 |
| CGH.L | 0.02690294 | -0.0350432 |
| CGH.R | 0.02991525 | -0.0199806 |
| CST.L | 0.01373639 | -0.0702227 |
| CST.R | 0.01481316 | -0.0799149 |
| EC.L | 0.03394982 | -0.0236498 |
| EC.R | 0.03302652 | -0.0184833 |
| FX_ST.L | 0.02943447 | 0.01486049 |
| FX_ST.R | 0.02415942 | -0.0194553 |
| IFO.L | 0.02982459 | -0.0335055 |
| IFO.R | 0.02293945 | -0.0582789 |
| PCR.L | 0.03321934 | 0.02940289 |
| PCR.R | 0.03359357 | 0.02793775 |
| PLIC.L | 0.02703531 | -0.0447676 |
| PLIC.R | 0.02303451 | -0.04982 |
| PTR.L | 0.03075398 | 0.05020077 |
| PTR.R | 0.0310768 | 0.04881825 |
| RLIC.L | 0.03119436 | 0.02555341 |
| RLIC.R | 0.02648144 | 5.96E-05 |
| SCR.L | 0.03445291 | 0.00972615 |
| SCR.R | 0.03500407 | 0.00937605 |
| SFO.L | 0.03111714 | 0.00856621 |
| SFO.R | 0.03095532 | 0.00184453 |
| SLF.L | 0.03432772 | 0.02459705 |
| SLF.R | 0.03541302 | 0.01758577 |
| SS.L | 0.03144969 | 0.03718058 |
| SS.R | 0.03234395 | 0.03191633 |
| UNC.L | 0.01796625 | 0.04371276 |
| UNC.R | 0.0243558 | 0.05270907 |

**Supplementary Table S10. The tract-based loading coefficients of the mode of diffusivity (MO) for the first two principal components (PCs).**

| **ID** | **PC1** | **PC2** |
| --- | --- | --- |
| ACR.L | 0.03166426 | -0.0369213 |
| ACR.R | 0.03393937 | -0.0329639 |
| ALIC.L | 0.0301057 | 0.05076734 |
| ALIC.R | 0.02515983 | 0.05208769 |
| CGC.L | 0.03957149 | -0.0420756 |
| CGC.R | 0.02974344 | -0.0544421 |
| CGH.L | 0.03839167 | 0.00032616 |
| CGH.R | 0.04402916 | 0.01139138 |
| CST.L | 0.01446464 | -0.0259607 |
| CST.R | 0.0094239 | -0.0245209 |
| EC.L | 0.02217057 | -0.0417693 |
| EC.R | 0.01878964 | -0.0336463 |
| FX_ST.L | 0.03893409 | -0.0482729 |
| FX_ST.R | 0.02452118 | -0.0128466 |
| IFO.L | 0.02562975 | -0.0140126 |
| IFO.R | 0.01948406 | -0.0158438 |
| PCR.L | 0.03669764 | 0.01974755 |
| PCR.R | 0.02654663 | 0.03037681 |
| PLIC.L | 0.03762616 | 0.05556387 |
| PLIC.R | 0.04240438 | 0.06219504 |
| PTR.L | 0.04004081 | -0.0417281 |
| PTR.R | 0.01752543 | -0.033538 |
| RLIC.L | 0.05103402 | -0.0195368 |
| RLIC.R | 0.04763034 | 0.0007211 |
| SCR.L | 0.04068907 | 0.04578894 |
| SCR.R | 0.04317396 | 0.02668904 |
| SFO.L | 0.03335869 | 0.05768729 |
| SFO.R | 0.02505951 | 0.02882416 |
| SLF.L | 0.03931724 | -0.013684 |
| SLF.R | 0.02307095 | 0.0032123 |
| SS.L | 0.0308718 | -0.0210945 |
| SS.R | 0.00765757 | -0.0196224 |
| UNC.L | 0.00936648 | -0.0204346 |
| UNC.R | 0.00190654 | 0.0017069 |

**Supplementary Table S11. Analysis overview, feature sets, sample sizes, and validation procedures.**

| **Analysis** | **Inputs** | **Analytic N** | **Adjustment/validation** |
| --- | --- | --- | --- |
| Structural MWAS | Global measures; 3 PCs each for area, thickness, volume | 200 children; 424 sessions | Mixed models; prespecified covariates; BH FDR p-value adjustment |
| Diffusion MWAS | 2 PCs each for FA, MD, MO | 120 children; 176 sessions | Mixed models; prespecified covariates; BH FDR p-value adjustment |
| Functional MWAS | Connectivity in 7 resting-state networks | 196 children; 400 sessions | Mixed models; prespecified covariates; BH FDR p-value adjustment |
| Sensitivity MWAS (30 days; h = 1 month) | All primary MWAS feature sets and structural, diffusion, and functional phenotypes | Structural: 200 children, 424 sessions; diffusion: 120 children, 176 sessions; functional: 196 children, 400 sessions | Closest same-child pair with \|stool–MRI interval\| ≤ 30 days; signed stool-minus-MRI age difference added to the prespecified covariates; mixed models; BH FDR p-value adjustment |
| Sensitivity MWAS (20 days; h = 2/3 month) | All primary MWAS feature sets and structural, diffusion, and functional phenotypes | Structural: 176 children, 327 sessions; diffusion: 101 children, 138 sessions; functional: 171 children, 316 sessions | Closest same-child pair with \|stool–MRI interval\| ≤ 20 days; signed stool-minus-MRI age difference added to the prespecified covariates; mixed models; BH FDR p-value adjustment |
| Sensitivity MWAS (15 days; h = 0.5 month) | All primary MWAS feature sets and structural, diffusion, and functional phenotypes | Structural: 162 children, 273 sessions; diffusion: 92 children, 120 sessions; functional: 153 children, 262 sessions | Closest same-child pair with \|stool–MRI interval\| ≤ 15 days; signed stool-minus-MRI age difference added to the prespecified covariates; mixed models; BH FDR p-value adjustment |
| Phenotype covarying patterns | Five microbiome blocks (GBM, GMM, species, MetaCyc pathways, and diversity) × four neuroimaging blocks (regional structural phenotypes, 9 structural PCs, 6 diffusion PCs, and 7 network-connectivity measures) | 26–102 children per tested feature pair after 5-MAD filtering (31–102 before filtering) | Covariate-adjusted participant-specific age slopes; Pearson correlations; 5-MAD filtering; BH FDR within each microbiome–neuroimaging block pair |
| rCCA | 78 microbial + 22 brain features | 107 children with complete multimodal data | 1000 permutations and repeated 75/25 train/test splits |
| Sparse canonical PLS | Same 78 microbial + 22 brain features as rCCA | 107 children with complete multimodal data (same cohort as rCCA) | 1000 permutations and repeated 90/10 train/test splits; score and feature-pattern concordance with the fixed CCA axes |
| CCA covarying patterns | Longitudinal microbiome and neuroimaging CCA1 and CCA2 scores | 29 children with at least two ages in both domains (CCA1 and CCA2) | Covariate-adjusted participant-specific age slopes; Pearson correlations; 5-MAD filtering; BH FDR across the two CCA modes |
| CCA cross-lag associations | Longitudinal microbiome and neuroimaging CCA1 and CCA2 scores; six prespecified directional lag windows (1–3, 3–6, and 6–12 months) | Retained window pairs: 44–75 children and 61–102 pairs; estimable autoregressive models: 20–69 children and 26–96 complete pairs | One-to-one within-child visit matching; adjustment for the prior outcome, exact lag, age, prespecified covariates, sequencing depth, and total brain volume; random-intercept models or participant-clustered HC1 SEs; 5-MAD filtering; BH FDR within each CCA mode |

Supplementary Text

Inclusion criteria of the study

Participants were recruited through research registries at both universities, local newborn nurseries, institutional centers focused on early brain development, community flyers, and university listservs. UNC/UMN BCP inclusion criteria were: (1) gestational age at birth of 37–42 weeks; (2) birth weight appropriate for gestational age; and (3) no major maternal pregnancy or delivery complications. Exclusion criteria were: (1) adoption; (2) diagnosis of autism, intellectual disability, schizophrenia, or bipolar disorder; (3) birth weight <2 kg; (4) neonatal hypoxia, defined as a 10-min Apgar score <5; (5) illness requiring >2 days of neonatal intensive care; (6) chromosomal or major congenital abnormalities; (7) prior abnormal MRI findings; (8) significant developmental delay or medical illness; (9) contraindication to MRI; (10) maternal pre-eclampsia, HIV positivity, or placental abruption; and (11) maternal alcohol or illicit drug use during pregnancy. For the present study, children also had to be younger than 3 years and have a fecal sample collected within 30 days of the corresponding MRI visit.

Fecal sample collection and processing

Parents were asked to collect a fecal sample from their child's diaper at home using the OMNIgene GUT kit (DNA Genotek, Ontario, Canada) within 24 hours of the scheduled MRI scan and bring the sample to the study visit. If a sample was not collected before the visit, it was collected on site when possible; otherwise, families were provided a pre-addressed envelope for home collection and return after the visit. Samples were processed within one week of collection. The material was loosened in a dry bead bath, transferred with a sterile pipette into sterile Eppendorf tubes, divided evenly between two tubes, and stored at −80 °C. Samples were then shipped to CosmosID, Inc. (Germantown, MD, USA) for analysis.

DNA extraction, library preparation and sequencing

All DNA extraction, library preparation, and sequencing were performed by CosmosID, Inc. (Germantown, MD, USA; now Cmbio), a CLIA-certified, ICH-GCP-compliant laboratory, using its standard deep-shotgun metagenomics workflow. Extracted DNA was quantified with a Qubit 4 fluorometer and Qubit dsDNA HS Assay Kit. Libraries were prepared with the Nextera XT DNA Library Preparation Kit (Illumina, San Diego, CA, USA) and IDT Unique Dual Indexes using 1 ng DNA input. DNA was fragmented with the Nextera XT fragmentation enzyme, unique dual indexes were added, and libraries were amplified for 12 PCR cycles. Libraries were purified with AMPure magnetic beads (Beckman Coulter, Brea, CA, USA), eluted in QIAGEN EB buffer, quantified by Qubit, and sequenced on an Illumina NovaSeq 6000 System with an S4 Flow Cell. After duplicate removal, adapter/PhiX removal, host depletion, and quality filtering, retained samples contained 3–118 million reads (mean, 17.9 million); 16 samples with fewer than 3 million post-QC reads were excluded.

MRI acquisition

All MRI was acquired without sedation during natural sleep. Infants were fed and swaddled, lighting was controlled, white noise was used, and a trained coordinator monitored the sleeping child throughout the session. In this protocol, ‘resting-state’ fMRI denotes spontaneous BOLD acquisition during monitored natural sleep rather than wakeful task-free rest. Imaging used Siemens 3 T Prisma scanners with a 32-channel phased-array coil. T1-weighted images used TR = 2,400 ms, TE = 2.24 ms, and 0.8-mm isotropic resolution; T2-weighted images used TR = 3,200 ms, TE = 564 ms, and 0.8-mm isotropic resolution. T2*-weighted rs-fMRI used TR = 800 ms, TE = 37 ms, 2-mm isotropic resolution, and 420 volumes. dMRI used 1.5-mm isotropic resolution, FOV = 210 × 210 mm, TE = 88 ms, TR = 2,365 ms, and multiband factor 5. Six b-values (500, 1,000, 1,500, 2,000, 2,500, and 3,000 s/mm²) used 9, 12, 17, 24, 34, and 48 directions, respectively; six interleaved b = 0 images were acquired.

Associations with global infant microbiome composition assessed by PERMANOVA

PERMANOVA identified age as the largest measured correlate of infant gut microbial beta diversity, accounting for 10.66% of variation in community composition (R² = 0.10661, p < 0.0001). Site and sex each accounted for approximately 0.35% of variation (R² ≈ 0.00348, p < 0.01), and birth weight and birth length each accounted for approximately 0.3% (R² ≈ 0.003, p < 0.05). Delivery mode accounted for 1.0% of variation (R² = 0.01000, p < 0.0001). Maternal education and feeding practice at four months each accounted for approximately 1.5% (R² ≈ 0.015, p < 0.0001). These estimates describe associations with microbial community composition and should not be interpreted as causal effects.

Sequencing depth was also associated with microbial composition (R² = 0.0030, p < 0.05). Gestational age did not meet the conventional significance threshold (R² = 0.0025, p = 0.052).

The joint PERMANOVA model explained 16.7% of microbial-community variation (R² = 0.167, p < 0.0001). The remaining variation may reflect unmeasured time-varying breastfeeding status, detailed diet, recent antibiotic exposure, intercurrent illness or gastrointestinal symptoms, environmental exposures, host genetics, and other factors. These variables were not available in this dataset and remain potential sources of residual confounding.

Principal-component interpretation of regional structural and diffusion measures

Because regional cortical structural measures and tract-based diffusion measures were high dimensional relative to the global anatomical and functional-network phenotypes, we used principal component analysis (PCA) for dimension reduction. Figure 4A shows the 30 regions with the largest absolute loadings for the first three PCs of cortical volume, surface area, and thickness; Figure 4B shows loadings across all white-matter tracts. Cortical volume and surface-area PCs showed similar spatial loading patterns. PC1 was dominated by positive loadings and PC2 by negative loadings, whereas PC3 included both positive and negative loadings. The largest absolute loadings were concentrated in orbital, insular, and cingulate regions for PC1; inferior and middle temporal, posterior cingulate, middle frontal, and right parahippocampal regions for PC2; and lateral temporal and orbitofrontal regions for PC3. Cortical-thickness PCs showed a different pattern: PC1 was predominantly negative, whereas PC2 and PC3 contained mixed positive and negative loadings. The largest absolute thickness loadings occurred in superior frontal and inferior parietal regions for PC1; cingulate, gyrus rectus, parahippocampal, and central regions for PC2; and orbital, insular, and occipitotemporal regions for PC3. Supplementary Tables S5–S7 report the corresponding regional loadings, and Supplementary Figure S5 shows loadings across all 148 cortical regions.

The diffusion PCs also showed distinct tract-loading patterns (Figure 4B; Supplementary Tables S8–S10). For FA, MD, and MO, PC1 included large loadings in major tracts such as the external capsule, superior longitudinal fasciculus, and superior corona radiata. PC2 differed across diffusion measures: FA emphasized the uncinate fasciculus and anterior corona radiata; MD emphasized the corticospinal tracts and inferior fronto-occipital fasciculus; and MO emphasized the posterior and anterior limbs of the internal capsule and the cingulum. These loading patterns were used to interpret the diffusion PCs in subsequent analyses.

Exploratory microbiome–neuroimaging associations (0.05 ≤ q < 0.25)

Supplementary Figures S6 and S8 and Supplementary Excel Table S1 display the complete set of associations with q < 0.25. Thirty-nine associations fell in the exploratory range 0.05 ≤ q < 0.25; the five associations with q < 0.05 are reported in the main text. Across the full q < 0.25 set, 84.1% involved morphological measures, 11.4% involved diffusion measures, and 4.5% involved functional networks. The remainder of this section discusses only the exploratory 0.05 ≤ q < 0.25 associations.

Among the exploratory taxonomic findings (Supplementary Figure S6), *Asaccharobacter celatus* and the genus *Asaccharobacter* were negatively associated with principal components of cortical area and volume. *Asaccharobacter* belongs to Eggerthellaceae and can convert dietary isoflavones to equol. Prior studies have linked equol-related exposures to both potentially beneficial and adverse outcomes in older populations, and one study reported greater *A. celatus* abundance in children older than 18 months with higher cognitive scores. These observations provide biological background but do not establish a mechanism for the exploratory associations in this infant cohort.

The exploratory pathway findings involved nucleotide, tricarboxylic-acid-cycle, carbohydrate (including trehalose and GABA degradation), amino-acid, and lipid metabolism. Within nucleotide metabolism, de novo purine biosynthetic pathways PWY0-162, PWY-2941, and P23-PWY were inversely associated with the first principal component of mode of anisotropy (MO; q = 0.10–0.25), which loaded most strongly on the internal capsule, cingulum, and superior corona radiata (Figure 4B). MO reflects diffusion-tensor shape and increased with age in this cohort. The contrast between these exploratory de novo purine associations and the q < 0.05 positive associations of purine-recycling pathways with total brain volume is hypothesis-generating and requires independent confirmation.

The genus *Gordonibacter* was positively associated with executive-control network connectivity (q = 0.078), in the same direction as the q < 0.05 species-level association for *Gordonibacter pamelaeae* shown in Figure 5 and Supplementary Figure S7. *Gordonibacter* species can metabolize ellagic acid to urolithins, which have been studied in relation to mitochondrial function, mitophagy, and inflammatory pathways. The genus-level result remains exploratory and does not establish that *Gordonibacter* or its metabolites alter infant functional connectivity. Associations with diffusion and functional phenotypes were less frequent than associations with morphology.

MWAS sensitivity analysis

To assess sensitivity to stool–MRI timing, we repeated all MWAS using the closest same-child stool and neuroimaging observations within maximum matching windows of 15 days (h = 0.5 month), 20 days (h = 2/3 month), and 30 days (h = 1 month). Each model retained the original prespecified covariates and additionally adjusted for the signed stool-minus-MRI age difference (microbiome age minus neuroimaging age, in months). Analytic sample sizes decreased with narrower windows: structural MWAS included 424, 327, and 273 sessions from 200, 176, and 162 children; diffusion MWAS included 176, 138, and 120 sessions from 120, 101, and 92 children; and functional MWAS included 400, 316, and 262 sessions from 196, 171, and 153 children for the 30-, 20-, and 15-day windows, respectively (Supplementary Table S11).

Seven associations met q < 0.05 in at least one sensitivity analysis, although the significant set differed across matching windows (Supplementary Figure S14A). In contrast, effect estimates for the five original MWAS signals changed little across windows (Supplementary Figure S14B): coefficient ranges were 0.209–0.226 for PWY0-1297 with total brain volume, 0.129–0.138 for PWY66-409 with total brain volume, −0.720 to −0.667 and −0.708 to −0.652 for *Actinomyces* sp. HMSC035G02 with cortical area PC3 and cortical volume PC3, respectively, and 0.529–0.548 for *Gordonibacter pamelaeae* with executive-control connectivity. The PWY0-1297 purine deoxyribonucleoside degradation association with total brain volume remained FDR-significant under the 30-day (q = 0.001) and 20-day (q = 0.007) windows; the similar 15-day estimate did not pass FDR correction (q = 0.067). Stable effect estimates with loss of FDR significance as sample size decreased are consistent with reduced precision under narrower matching windows, although other window-related changes cannot be excluded.
